# Cholesterol inhibits assembly and activation of the EphA2 receptor

**DOI:** 10.1101/2024.06.10.598255

**Authors:** Ryan J Schuck, Alyssa E Ward, Amita R Sahoo, Jennifer A Rybak, Robert J Pyron, Thomas N Trybala, Timothy B Simmons, Joshua A Baccile, Ioannis Sgouralis, Matthias Buck, Rajan Lamichhane, Francisco N Barrera

## Abstract

The receptor tyrosine kinase EphA2 drives cancer malignancy by facilitating metastasis. EphA2 can be found in different self-assembly states: as a monomer, dimer, and oligomer. However, our understanding remains limited regarding which EphA2 state is responsible for driving pro-metastatic signaling. To address this limitation, we have developed SiMPull-POP, a single-molecule method for accurate quantification of membrane protein self-assembly. Our experiments revealed that a reduction of plasma membrane cholesterol strongly promoted EphA2 self-assembly. Indeed, low cholesterol caused a similar effect to the EphA2 ligand ephrinA1-Fc. These results indicate that cholesterol inhibits EphA2 assembly. Phosphorylation studies in different cell lines revealed that low cholesterol increased phospho-serine levels, the signature of oncogenic signaling. Investigation of the mechanism that cholesterol uses to inhibit the assembly and activity of EphA2 indicate an in-trans effect, where EphA2 is phosphorylated by protein kinase A downstream of beta-adrenergic receptor activity, which cholesterol also inhibits. Our study not only provides new mechanistic insights on EphA2 oncogenic function, but also suggests that cholesterol acts as a molecular safeguard mechanism that prevents uncontrolled self-assembly and activation of EphA2.

## Introduction

The receptor tyrosine kinase EphA2 is activated by binding to ephrin ligands on opposing cells. The establishment of cell-to-cell contacts through EphA2-ephrin initiates a signaling cascade that regulates cell morphology, adhesion, migration, and survival. Such processes are important for proper embryonic development, neuronal plasticity, wound healing, and homeostasis of adult epithelial tissues. However, misregulation of EphA2 signaling contributes to human disorders and pathological states, including cancer, and EphA2 is overexpressed in various cancer types, including breast, ovarian, prostate, and pancreatic tumors^1–5^.

EphA2 participates in two signaling modes: ligand-dependent and ligand-independent. The canonical ligand-dependent signaling occurs after activation of EphA2 by binding of its protein ligands, including ephrinA1. Ligand binding causes auto-phosphorylation of tyrosines as EphA2 dimerizes and forms higher-order oligomers and clusters^6–8^. Ligand-dependent signaling inhibits oncogenic phenotypes, as it is characterized by maintenance of physiological cell-to-cell contacts and a decrease in cell proliferation and migration^7–9^. However, EphA2 can also signal in the absence of ligand binding; ligand-independent EphA2 signaling is oncogenic and is characterized by increased phosphorylation of serines, including S897. EphA2 serine phosphorylation is carried out by major signaling axes such as cAMP/PKA, AKT/mTORC1, and RAS/ERK^7,10–12^. Inhibition of ligand-independent activation of EphA2 represents a potential target for cancer therapeutics. However, the mechanisms underlying EphA2 noncanonical signaling are poorly understood.

Membrane lipids often affect integral membrane protein conformation and activity, and these lipid effects can occur through direct (allosteric) binding or by indirect mechanisms^13,14^. The composition of the membrane also impacts the activity and spatial recruitment of intracellular interacting partners. As a result, defects in lipid metabolism are associated with various human diseases^15,16^. Therefore, it is necessary to understand protein-lipid interactions to fully establish the molecular basis of diseases caused by the malfunction of membrane proteins. Cholesterol (Chol) is the most abundant molecule in the plasma membrane of human cells, representing 30-40 % of all lipids^17^. Chol influences membrane structure characteristics such as fluidity, curvature, stiffness, and permeability^17–19^. Additionally, Chol impacts protein-protein interactions, enzyme activity, signal transduction, and intracellular trafficking^18,19^. However, it is currently unknown whether Chol impacts EphA2 oligomerization or activity.

In this work, we investigate the influence of Chol on EphA2 self-assembly at the single-molecule level and Chol’s impact on EphA2 activity. Our findings indicate a model where Chol negatively regulates EphA2 oligomerization and suppresses the oncogenic, ligand-independent signaling. We propose that the Chol-mediated inhibition of EphA2 oncogenic activation results from control of the cAMP/PKA signaling network.

## Results

### Single-molecule quantification of EphA2 oligomerization in a native-like membrane environment

EphA2 adopts different assembly states in the plasma membrane, as it can be found as a monomer, dimer, and oligomers that come together to form micro-sized clusters. It is important to understand the self-assembly of EphA2, as it determines its function. Current methods to study the oligomerization of membrane proteins frequently lack the ability to quantify the distribution of oligomeric states accurately. To address this limitation, we developed SiMPull-POP (Single-Molecule Pulldown - Polymeric-nanodisc Oligomer Photobleaching). SiMPull-POP is a single-molecule method that quantifies the oligomerization of membrane proteins. To apply SiMPull-POP to EphA2, we transfected HEK293T cells, which do not express detectable levels of endogenous EphA2 (**Figure S1**), with a plasmid coding for EphA2-GFP. We estimated that the EphA2-GFP density in the membrane had a median value of 253 molecules per square micron (**Figure S1**), similar to the physiological expression levels of EphA2^20–23^. HEK293T membrane fractions were solubilized with the copolymer diisobutylene/maleic acid (DIBMA), which forms ∼25 nm lipid nanodiscs termed DIBMALPs (**Figure 1A**). DIBMALPs provide a more physiological reconstitution system than detergents and protein-based nanodiscs, as they retain a native-like lipid composition^24^. The formation of DIBMALPs was confirmed with negative-stain transmission electron microscopy (TEM) (**Figure S2**). At our low receptor density, the probability that two non-interacting receptors are randomly captured in a single DIBMALP is negligible^25^ (**Figure S1**). DIBMALPs containing EphA2-GFP were purified using single-molecule pulldown (SiMPull) on a microfluidic chamber^26^. For SiMPull-POP, we used quartz slides functionalized with a biotinylated EphA2 monoclonal antibody immobilized on the slide surface via NeutrAvidin (**Figure 1A**). The non-specific/unbound sample was washed away. We imaged our samples via total internal reflection fluorescence (TIRF) microscopy, which revealed individual DIBMALPs (**Figure 1B**). As a negative control, we repeated experiments without the EphA2 antibody immobilized on the slide.

**Figure 1:**
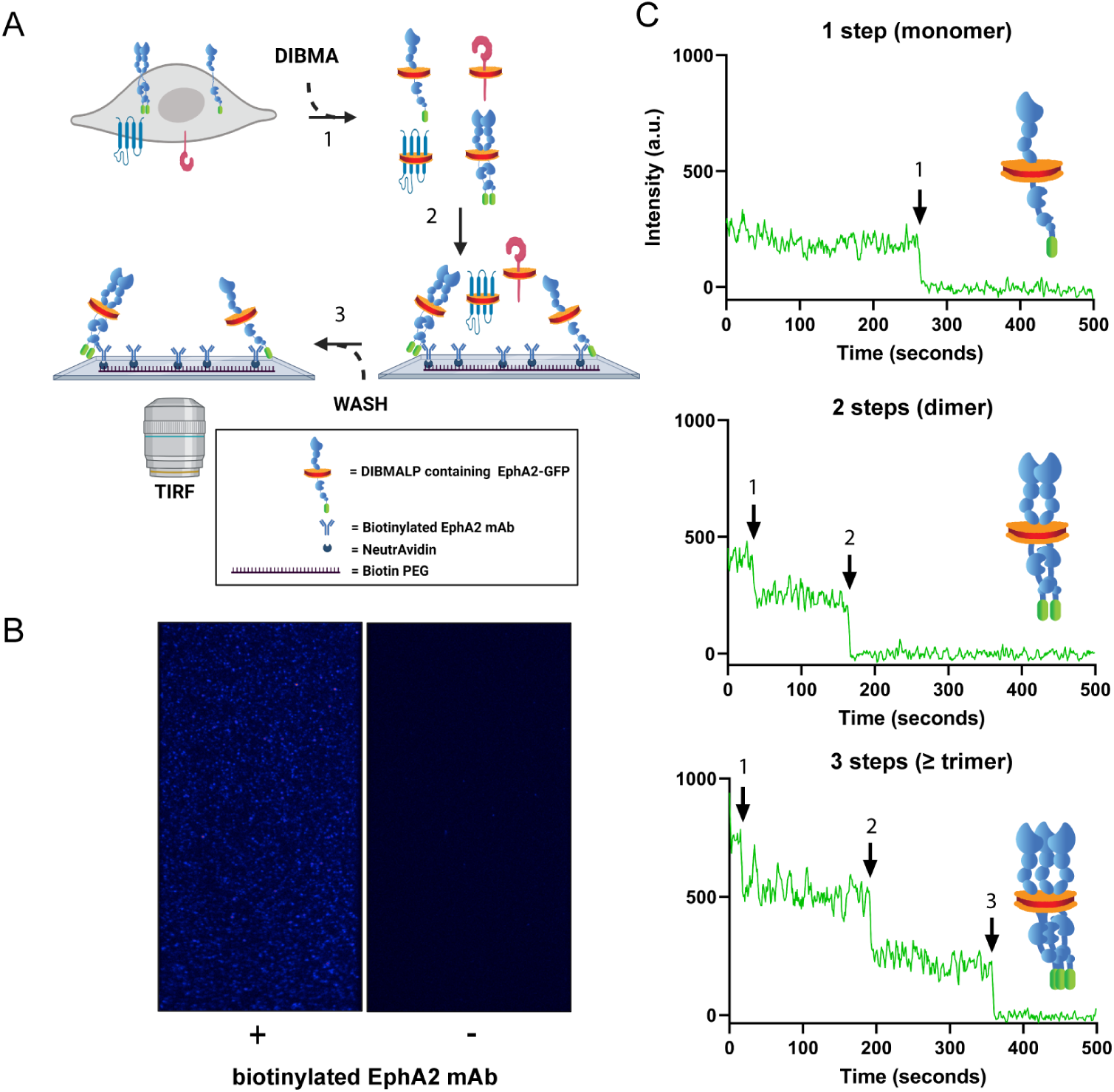
GFP photobleaching analysis via SiMPull-POP reports on EphA2 oligomerization in a native-like membrane environment. **(A)** Schematic representation of sample preparation and workflow for SiMPull-POP. 1-Membrane fractions containing EphA2-GFP were solubilized with the amphipathic copolymer DIBMA to generate DIBMALPs. 2-EphA2-GFP DIBMALPs were immobilized on a functionalized microscope slide displaying an EphA2 antibody. 3-DIBMALPs devoid of EphA2-GFP are washed away before imaging. **(B)** Representative single-molecule TIRF image in the presence (left) and absence (right) of EphA2 antibody. Each blue spot represents a DIBMALP containing EphA2-GFP. **(C)** Representative GFP photobleaching traces showing a stepwise decrease in GFP intensity over time; arrows represent individual photobleaching events. Photobleaching steps are used to infer EphA2 oligomerization status.

In these conditions, GFP fluorescence was negligible (**Figure A1 and S3**), demonstrating the success of the SiMPull approach to inform specifically for EphA2-GFP.

We next applied the single molecule photobleaching step analysis that we and others had previously developed in polymeric nanodiscs^27–29^. The resulting SiMPull-POP protocol analyzes the fluorescence of individual DIBMALPs over time and identifies GFP photobleaching steps, which are used to infer the oligomeric status of EphA2. We observed that EphA2-GFP DIBMALPs exhibited photobleaching events characterized mainly by one and two steps (**Figure 1C**). We also detected fewer traces with three or more photobleaching steps. These were collectively binned as higher-order oligomers (**Figure 1C**), as reported elsewhere^27,30^. These results show that SiMPull-POP captures and resolves EphA2 oligomeric states in the absence of exogenous ligands, which is in agreement with prior observations^31^.

### SiMPull-POP captures ligand-induced FKBP dimerization

To establish the robustness of SiMPull-POP, before performing a quantitative analysis of the EphA2 data, we applied the method to a well-described dimerization-inducible system. We studied the self-assembly of the FK506 binding protein (FKBP), which dimerizes upon binding to the AP ligand. GFP-tagged FKBP was immobilized at the membrane *via* myristoylation (Myr-FKBP-GFP). We applied SiMPull-POP to HEK293T cells expressing Myr-FKBP-GFP, under both control conditions or treated with the AP ligand (**Figure 2A**). DIBMALPs capturing Myr-FKBP-GFP were isolated on the slide surface using a biotinylated GFP antibody, and any non-specifically bound sample was efficiently washed away (**Figure S4**). As expected, under control conditions, the large majority of Myr-FKBP-GFP exhibited photobleaching characterized by a single step, as shown in **Figure 2B**. The addition of the AP ligand caused a significant decrease in one-step photobleaching and a concomitant increase in two-step photobleaching. These data suggest that SiMPull-POP allows to quantitatively tracking changes in oligomerization.

**Figure 2:**
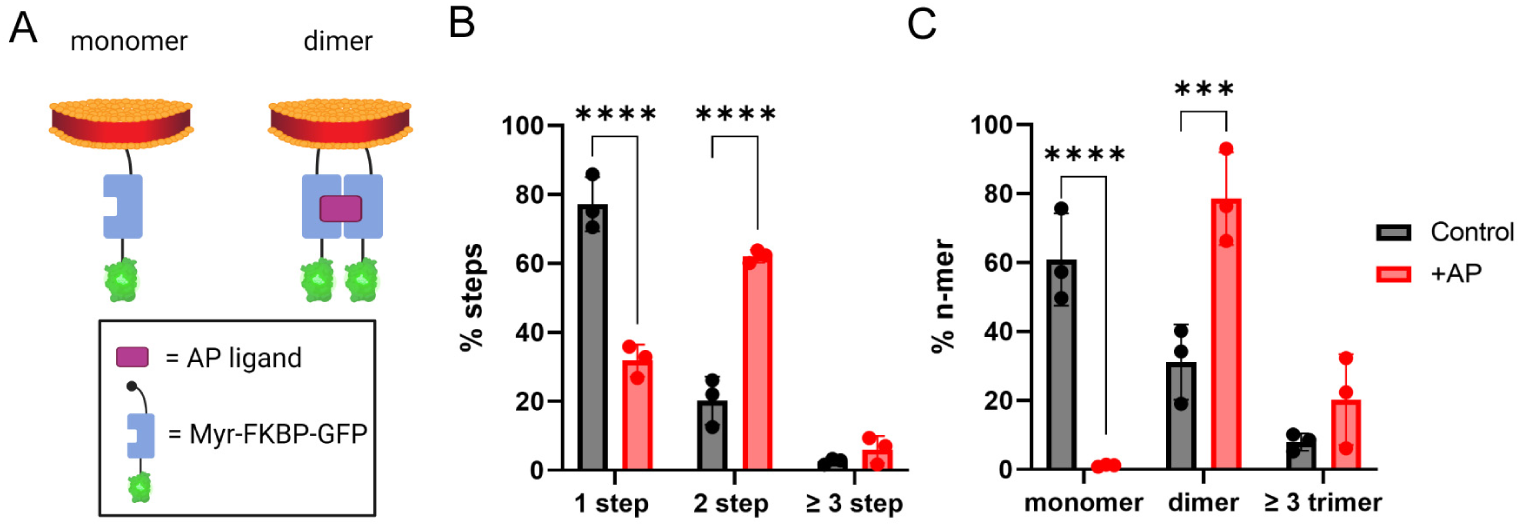
SiMPull-POP reports on FKBP dimerization. **(A)** Schematic representation of DIBMALPs containing Myr-FKBP-GFP (monomer, left). FKBP-GFP dimerization is induced by the AP ligand (dimer, right). **(B)** Experimental step distribution of FKBP-GFP photobleaching in control conditions (black) and in the presence of AP ligand (red). **(C)** Calculated oligomeric distribution corrected for 70% maturation efficiency of GFP. *p*-values are from two-way ANOVA followed by Tukey multiple comparison test, ***, *p* ≤ 0.001; ****, *p* ≤ 0.0001.

It is important to consider that a photobleaching step distribution does not directly inform the oligomerization status of GFP-labeled proteins^25^. Since GFP has a maturation efficiency of ∼70% in the cell^32,33^, a significant level of FKBP-GFP dimers will carry one normal GFP plus a non-fluorescent immature copy. Therefore, incomplete GFP maturation causes the photobleaching step data to underestimate dimers and higher-order oligomers while overestimating monomers. To account for the GFP maturation efficiency, we devised a theoretical probability distribution that allows us to correct the photobleaching step data, as described in the Methods section. The theoretical probability distribution allows us to extract an accurate distribution between monomers, dimers, and oligomers from the raw photobleaching step data (**Figure 2C**). After the GFP maturation correction, the data showed a larger population of oligomers, as expected. In agreement with our initial expectations, in the presence of ligand, no monomer was observed and most (∼80%) of Myr-FKBP-GFP was found as a dimer, while in the absence of ligand, the monomer was the most abundant state. These data indicate that our approach effectively captures ligand-induced dimerization in a native-like membrane environment. Furthermore, the results validated the use of SiMPull-POP to investigate the oligomerization of membrane proteins.

### Cholesterol reduction promotes oligomerization of EphA2

After benchmarking SiMPull-POP, we applied it to study physiological factors that control the self-assembly of EphA2. Quantification of photobleaching data showed that, in control conditions, the most abundant EphA2 state was the monomer, with lower levels of dimers and oligomers (≥ trimer) (**Figure 3 A-C**). We next treated samples, prior to DIBMALP formation, with the ligand EphrinA1-Fc (EA1), which causes EphA2 clustering. ^3,34,35–37^ We observed that addition of EA1 increased EphA2 self-assembly, as expected. Specifically, there was a significant reduction in the percentage of monomers and a large increase in oligomers (**Figure 3C**). These results demonstrate that SiMPull-POP effectively detects the EphA2 clustering induced by EA1 within the cellular plasma membrane^3,34^.

**Figure 3:**
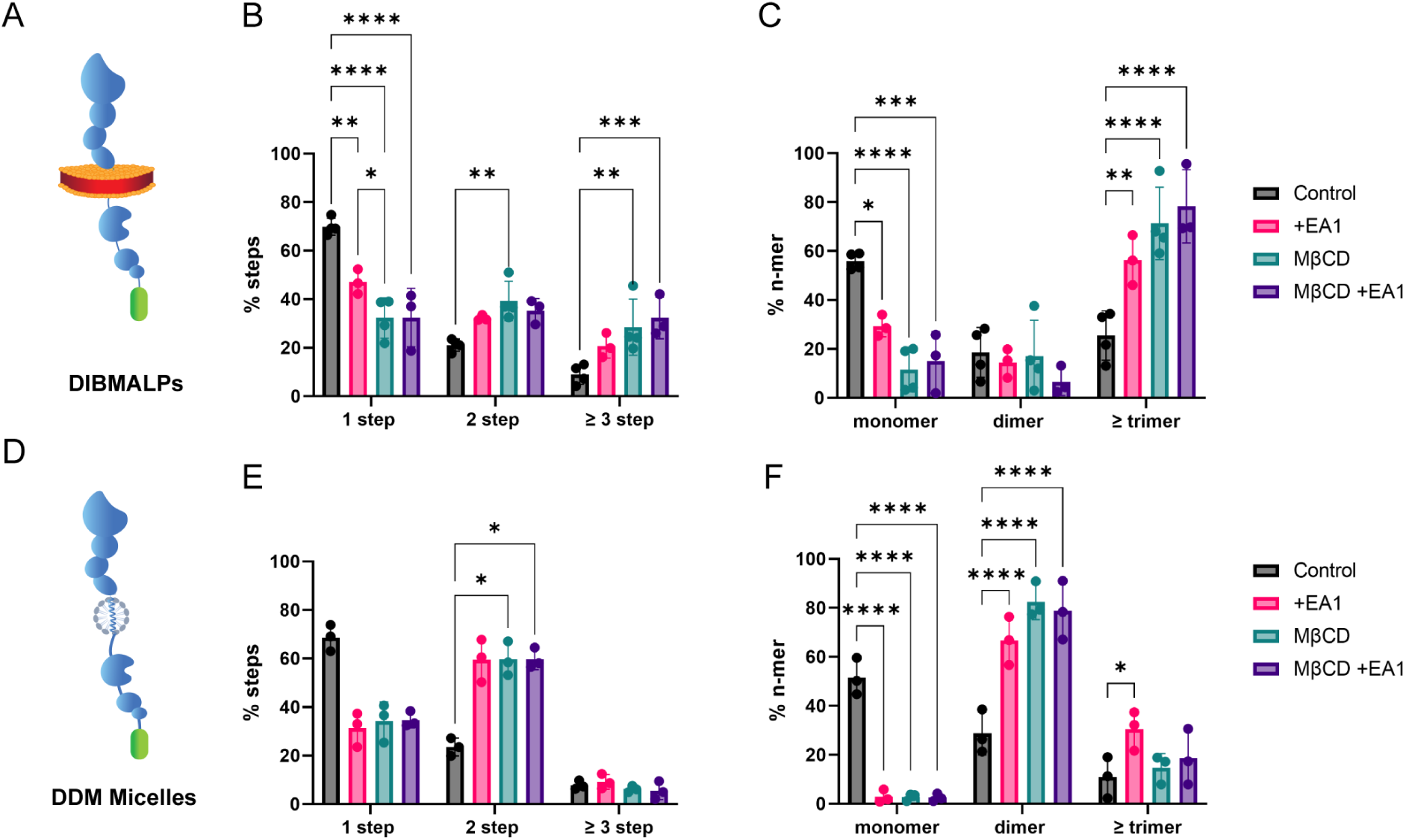
Cholesterol reduction promotes EphA2 oligomerization in the absence of ligand. **(A)** Schematic of DIBMALP containing an EphA2-GFP monomer. **(B)** Step distribution of control DIBMALPs (black) or those formed from cells treated with EA1 (pink), MβCD (blue) or both (magenta). **(C)** Oligomeric distribution calculated from data in panel B. **(D)** Schematic representing DDM micelles containing EphA2-GFP. **(E)** Step distribution of DDM-solubilized EphA2-GFP in the same conditions as in DIBMALPs. **(F)** Oligomeric distribution of DDM-solubilized EphA2-GFP photobleaching data. *p*-values are from two-way ANOVA followed by Tukey multiple comparison test.*, *p* ≤ 0.05; **, *p* ≤ 0.01; ***, *p* ≤ 0.001; ****, *p* ≤ 0.0001.

Next, we studied if changes in the lipid environment impact EphA2 self-assembly. We were interested in understanding if cholesterol (Chol) affects the self-assembly of EphA2. In order to answer this question, HEK293T cells expressing EphA2-GFP were treated with methyl-β-cyclodextrin (MβCD) to lower Chol levels^38–40^. We verified that MβCD treatment led to a significant reduction (∼40%) in Chol without impacting cell viability (**Figure S5A-B**).

We observed that Chol removal caused a large reduction in monomers and a corresponding increase in oligomers, which became the predominant population (**Figure 3C**). These data indicate that a reduction in plasma membrane Chol promotes EphA2 oligomerization. A comparison between the effects of MβCD and EA1 revealed striking similarities, as both treatments induced a large decrease in monomers and an increase of EphA2 oligomers. Indeed, DIBMALPs generated in the presence of both MβCD and EA1 showed similar results to MβCD treatment alone.

To rule out that MβCD could cause an artifact in DIBMALP formation, we applied this treatment to Myr-FKBP-GFP. We expected Chol to cause no effect on this control protein since the dimerization motif is outside the plasma membrane. **Figure S6** shows that, indeed, MβCD had no effect on Myr-FKBP-GFP dimerization, supporting that the self-assembly changes that we observed are specific for EphA2. Taken together, our data led us to hypothesize that the high levels of Chol in the plasma membrane ensure that EphA2 clustering does not occur in the absence of ligand. This action might serve as a potential safeguard mechanism that prevents non-specific EphA2 activation and constitutes a new physiological role for Chol.

As a control for the oligomerization changes observed in DIBMALPs, we quantified the oligomeric status of EphA2 after solubilization with the detergent dodecyl-β-maltoside (DDM). We expected that solubilizing EphA2 into DDM micelles would destabilize the formation of oligomers, as observed for other membrane complexes^41^. Indeed, we found that DDM micelles primarily captured EphA2-GFP monomers and dimers, as we observed significantly reduced oligomer levels (**Figure 3F**). These results suggest that detergent treatment interferes with the study of EphA2 oligomers. Nevertheless, in alignment with the results in DIBMALPs, we observed that EphA2-GFP in micelles displayed increased self-assembly upon Chol reduction. Taken together, our data show that a reduction of Chol promotes EphA2 oligomerization. These results indicate that Chol is an inhibitor of EphA2 self-assembly.

### Cholesterol reduction promotes ligand-independent EphA2 activity

Next, we investigated if the oligomeric changes induced by Chol led to a change in EphA2 activity^1–3,36,42,43^. We assessed the effect of MβCD treatment on EphA2 activity by western blots with phospho-specific EphA2 antibodies. We tracked ligand-independent (oncogenic) activation where serine kinases, like cAMP-activated protein kinase (PKA), phosphorylates S897 and other residues^44–46^. We also studied the phosphorylation of residue Y588 (pY588), which increases after ligand activation.

We first tested ligand activation in the HEK293T cells used for SiMPull-POP. We observed that EA1 (**Figure S7**) did not affect pS897 (**Figure 4A**), while it increased pY588 phosphorylation (**Figure S8**), as expected^34^. Interestingly, Chol depletion with MβCD did increase pS897 levels (**Figure 4A**). We repeated these experiments in the malignant melanoma A375 cell line, where EphA2 is endogenously expressed. Consistently, similar to the findings in HEK293T cells, the extraction of Chol *via* MβCD also led to increased pS897 levels (**Figure 4B**), while pY588 remained unchanged (**Figure S8**). We also observed this effect for the epidermal carcinoma cell line A431, which expresses higher levels of EphA2, where MβCD doubled phosphorylation at S897 (**Figure S8**). We next studied whether Chol changes affect ligand activation of EphA2. This was accomplished by performing a titration study to quantify the efficacy of EA1 in inducing Y588 phosphorylation. We observed that MβCD did not alter the effect of EA1 (**Figure S8**) in A375 cells, suggesting that Chol does not impact the ligand-dependent activation of EphA2. To ensure that the MβCD treatment specifically reduced the level of Chol but not other lipids, we performed lipidomics in A375 cells. The results confirmed that our MβCD treatment protocol did not significantly affect phospholipid or sphingomyelin levels (**Figure S9**). Altogether, our results show that Chol extraction induces a statistically significant increase in pS897 across different cell lines, which is the signature for oncogenic EphA2 ligand-independent activation.

**Figure 4:**
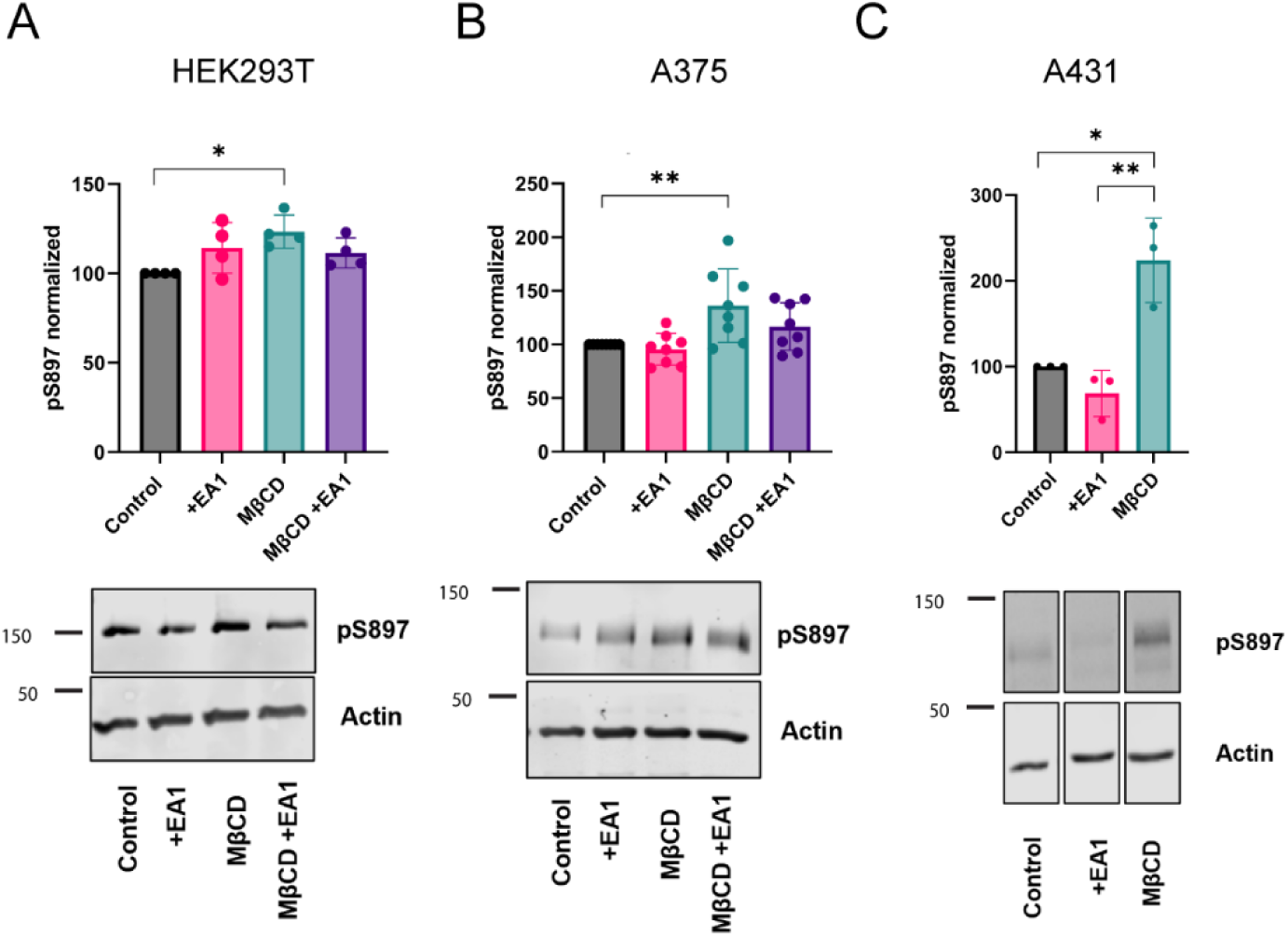
Extraction of cholesterol increases EphA2 Ser phosphorylation. Western blot analysis of EphA2 pS897 in HEK293T **(A)**, A375 **(B)** and A431 **(C)** cells. We show pS897 quantification (mean ± S.D) and representative blots. *p*-values from one-way ANOVA followed by Mann-Whitney *U* or *t* test. *, *p* ≤ 0.05; **, *p* ≤ 0.01.

Next, we used alternative means to lower cellular Chol levels. When we pharmacologically inhibited Chol synthesis with the reagent zaragozic acid^47^, we observed increased pS897 levels as well (**Figure S10**), in agreement with the MβCD results (**Figure 5**). Treatment with Zaragozic acid produced a ∼15% reduction in Chol content (**Figure S5E**), which was smaller than the ∼30% reduction caused by MβCD (**Figure S5C**). These results suggest that a moderate decrease in Chol levels is enough to change EphA2 activity. Taken together, the data obtained in HEK293T, A375, and A431 cells show that a reduction in Chol levels promotes ligand-independent activation of EphA2, which signals for oncogenic phenotypes.

**Figure 5:**
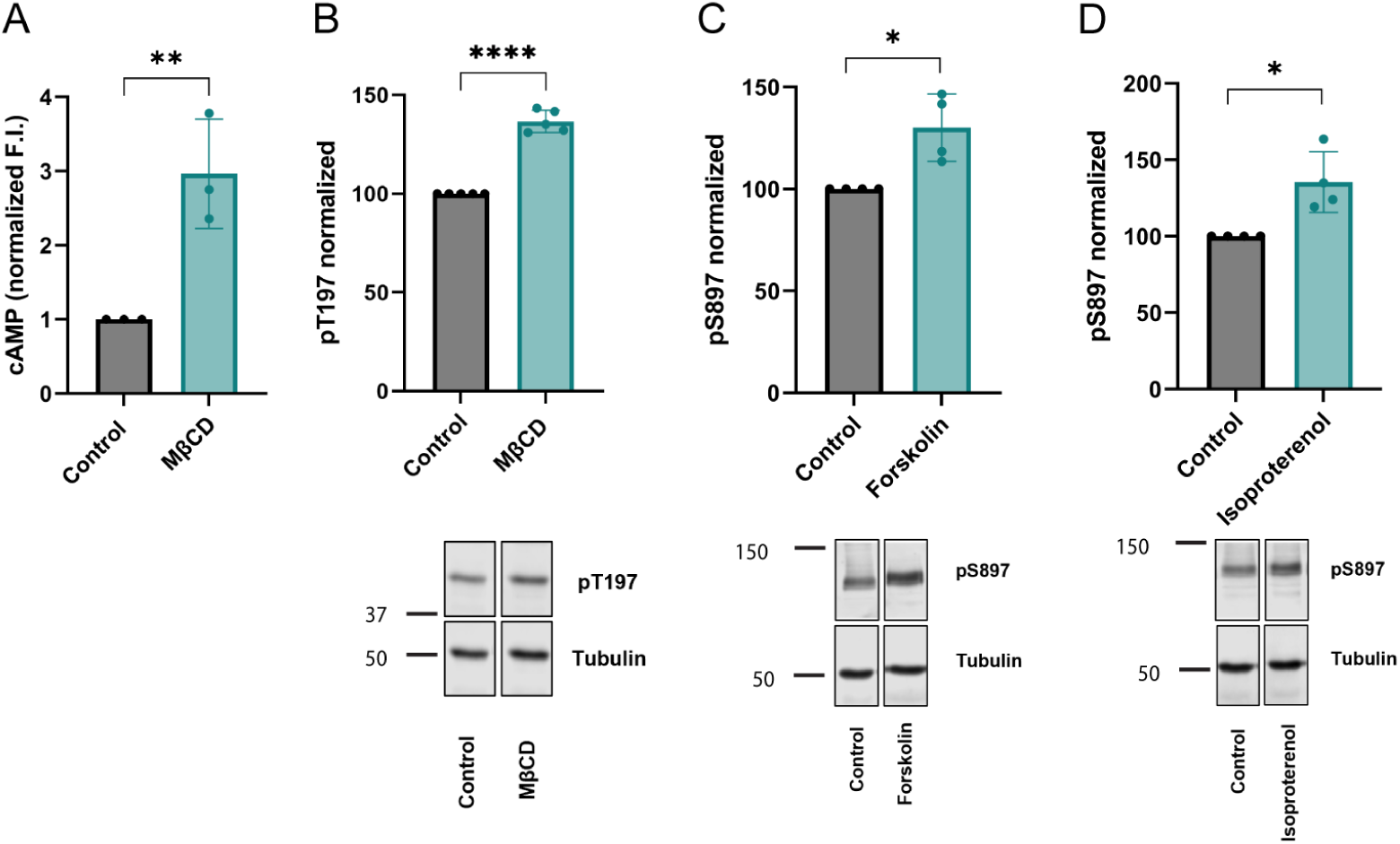
Activation of cAMP-dependent protein kinase and upstream β-AR promote pS897 EphA2. **(A)** cAMP quantification in HEK293T cells transduced with Red Up cADDis cAMP biosensor following treatment with MβCD. **(B)** Western blot analysis and quantification of PKA T197 phosphorylation in A375 cells following treatment with MβCD. **(C-D)** Western blot analysis and quantification of EphA2 S897 phosphorylation in A375 cells following treatment with forskolin and isoproterenol, respectively. Data shown in panel B are normalized to the respective total PKA signal in Figure S15A. Data shown in panels C-D are normalized to the respective total EphA2 signal in Figure S15B-C. Quantitative comparisons between treatments were made with respect to normalized control conditions. Bar graphs show mean ± S.D., *p*-values in panels A-D are from an unpaired t-test. *, *p* ≤ 0.05; **, *p* ≤ 0.01; ****, *p* ≤ 0.0001.

### Cholesterol does not regulate EphA2 in cis

What is the molecular mechanism that Chol uses to impact EphA2 activity? To answer this question, we first investigated whether the binding of Chol to the transmembrane (TM) domain of EphA2 affects self-assembly^34,46,48,49^. We used a disulfide crosslinking assay that we previously employed to demonstrate that the lipid phosphatidylinositol 4,5-bisphosphate regulates the dimerization of the TM of EphA2^27^. However, this assay suggested Chol does not alter the tendency of the TM of EphA2 to dimerize in the two synthetic lipid compositions assayed (**Figure S11**). Next, we performed molecular dynamics simulations. The microsecond-long simulations did not show a clear change in dimerization or binding of Chol to the TM of EphA2 (**Figure S12-S13 & Table S1**). These results suggest that Chol uses a mechanism other than acting as an *in cis*, allosteric ligand of EphA2.

We also investigated if changes in Chol levels altered the distribution of EphA2 in the plasma membrane. Laser scanning confocal microscopy experiments showed no noticeable effect on the cellular distribution of EphA2 upon MβCD treatment (**Figure S7**). We next performed an assay that determines the partitioning of molecules between Chol-enriched liquid-ordered (*L_o_*) domains and more fluid liquid disordered (*L_d_*) domains^50,51^. To this end, we formed phase-separated giant plasma membrane vesicles (GPMVs) derived from HeLa cells transfected with the EphA2-GFP plasmid^51^. These experiments indicated that EphA2-GFP strongly partitions to Chol-poor *L_d_* membrane regions regardless of MβCD treatment (**Figure S14**). The confocal and GPMV data suggest that Chol does not regulate EphA2 activity through large changes in plasma membrane distribution. Taken together, our data suggest that the conventional ways by which Chol affects membrane protein activity are not behind the EphA2 changes we observed. We therefore pivoted to consider a different hypothesis, which is that Chol acts *in trans* through other proteins.

### Cholesterol reduction leads to increased cAMP levels and enhances PKA activity

PKA is a major kinase that phosphorylates EphA2 serine residues^5,44,45^, and PKA function requires the secondary messenger cAMP^52,53^. It has been reported that Chol depletion can increase cAMP levels and activate PKA in different cell types^54–57^. We investigated if Chol reduction also caused cAMP increases in our experimental system. We tested the levels of cAMP in HEK293T cells using the cAMP biosensor cADDis. We observed that treatment with MβCD significantly increased cAMP levels (**Figure 5A**), suggesting that a drop in Chol levels could activate EphA2 *in trans via* PKA. To test this hypothesis, we determined if PKA activity also increased upon Chol depletion. We used western blot to track the activity of PKA, as reported by phosphorylation of the activity-dependent PKA residue T197. We indeed observed increased phosphorylation of PKA T197 after treatment with MβCD in A375 cells (**Figure 5B & S15**). We additionally employed an established pharmacological approach to increase cAMP levels, the adenylate cyclase activator forskolin^54,56,58^. We observed that forskolin treatment also led to higher pS897 EphA2 (**Figure 5C**). These results support the idea that Chol does not act directly on EphA2 but that it exerts its effect *in trans* through activation of PKA.

We investigated next if β-adrenergic receptors (β-AR), which are upstream of the cAMP enzyme adenylate cyclase, were activated by a drop in Chol levels. We activated βAR with Isoproterenol (Iso), an β_1_/β_2_-AR agonist that is an analog of epinephrine^55^. In agreement with this idea, isoproterenol treatment also promoted EphA2 phosphorylation at residues Ser897 (**Figure 5D**). Taken together, our data suggest that Chol reduction leads to a β-AR-mediated increase in cAMP, which activates PKA to phosphorylate EphA2 at serine residues.

## Discussion

Here, we report the development of the SiMPull-POP method. This new approach is able to uncover the oligomeric status of transmembrane and membrane-anchored proteins in the native-like lipid composition provided by DIBMALPs. Unlike other methods, SiMPull-POP quantifies the percentage of individual oligomeric species instead of providing an average value. The single-molecule resolution of SiMPull-POP allows it to work with minute amounts of sample (in the pM-nM range). SiMPull-POP revealed that Chol depletion promoted EphA2 self-assembly in the absence of added ligands. We observed that Chol reduction also increased Ser897 phosphorylation, which is an oncogenic signature of EphA2. Our results, therefore, identify Chol as a strong inhibitor of EphA2 assembly and ligand-independent activation.

We performed extensive studies to unravel the molecular mechanism by which Chol controls EphA2 activity and assembly. We first explored the simplest-case scenario, whereby Chol directly influences the TM region of EphA2 to alter its dimerization or membrane localization. However, computational, liposome and GPMV assay data argued against Chol directly affecting EphA2. Therefore, we turned our attention to PKA, which regulates EphA2 activity by phosphorylation and promotes the oncogenic phenotype.

We observed that Chol reduction increased cAMP levels and caused the concomitant PKA activation. Given these findings, we propose a model in which Chol depletion promotes oncogenic assembly and activity of EphA2 through activation of cAMP/PKA signaling. However, we cannot fully rule out the involvement of other serine kinases. Our data also indicate that cAMP/PKA activation results from β-AR activation caused by a decrease in Chol levels **(Figure 6)**. Previous reports have used experimental^59^ and computational approaches^60,61^ to show that Chol binds to the β_2_-AR. Chol has been recently identified as necessary for β_2_-AR dimerization^62^. Importantly, functional studies in cardiomyocytes have shown that Chol depletion activated β_2_-AR^63,64^, which is in agreement with our model. Our data, therefore, suggest that β−AR is a new regulator of EphA2.

**Figure 6:**
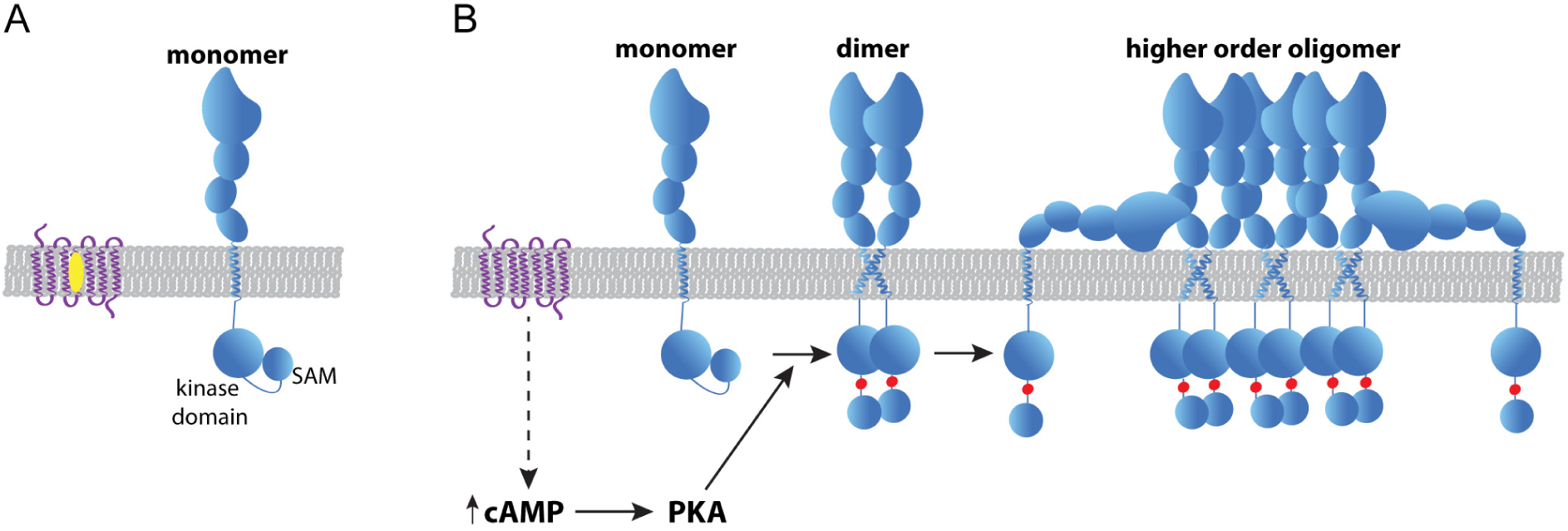
Proposed model of how Chol regulates EphA2 self-assembly & signaling. **(A)** In the presence of normal levels of Chol (yellow oval), EphA2 (blue) can be found as a monomer and displays low Ser897 phosphorylation. We propose that in this state the kinase domain interacts with the SAM domain. **(B)** When Chol content is reduced, β-AR (purple) activity increases, promoting cAMP/PKA signaling that enhances Ser897 phosphorylation (red dot). This forces the kinase-SAM linker into an open conformation, which promotes higher-order oligomers independent from ligand stimulation.

EphA2 clustering after ligand stimulation activates the kinase domain of the receptor and causes tyrosine auto-phosphorylation. Intriguingly, enhanced EphA2 self-assembly by Chol depletion does not increase pY588 in A375 and HEK293T cells. However, MβCD caused increased phosphorylation in Y588 in A431 cells. These results suggest that differences in the cellular context can modulate the physiological effect of Chol.

Our data indicate that Chol controls both EphA2 self-assembly and serine phosphorylation. It has been recently proposed that phosphorylation of S897 induces a conformational change in the linker region connecting the kinase and SAM domains, which results in enhanced EphA2 oligomerization^31,44,58^. In this hypothesis, the negative charges introduced by S897 phosphorylation block the interaction between the two domains of the same chain, which then adopt an extended conformation. In this state, the SAM and kinase domains are be able to dimerize with other parallel EphA2 chains, leading to increased self-assembly (**Figure 6**) compatible with the formation of EphA2 clusters^65^.

Cholesterol exerts direct and indirect effects on membrane-spanning and membrane-associated proteins. Such a variety of interactions can affect protein structure, activity, and localization within the plasma membranes^40,66–68^. Previous work with the epidermal growth factor receptor (EGFR) showed that Chol inhibits ligand-independent activation, as reduced Chol content promotes EGFR oligomerization and stimulates its pro-oncogenic activity^69–73^. However, the molecular mechanism behind how Chol regulates EGFR oligomerization and activity remains unknown. Here, we uncovered the mechanism that Chol uses to inhibit EphA2 assembly and oncogenic signaling through the β-AR/cAMP/PKA/EphA2 signaling axis. Our results highlight the anti-oncogenic effect of Chol, and suggest that this key lipid has a widespread inhibitory effect on receptor tyrosine kinases that drive tumor malignancy.

## Methods

### Plasmid constructs

Plasmid encoding full-length EphA2 containing a C-terminal turboGFP tag was obtained from Origene (Accession number: RG205725). Plasmid encoding Myr-FKBP-EGFP was a kind gift from Dr. Adam Smith (Texas Tech University).

### Cell culture and transfection

HEK293T, A375, A431, and HeLa cells were purchased from ATCC and maintained at 5% CO_2_ and 37 °C in Dulbecco’s Modified Eagle’s Medium (DMEM) supplemented with glucose, 10% fetal bovine serum (FBS), and 100 U/mL penicillin-streptomycin. Cells were passed at 80% confluency and were not used beyond 30 passages. Cell lines were tested for mycoplasma contamination via a PCR detection kit (Abcam) according to the manufacturer’s protocol. HEK293T and HeLa cells were transiently transfected with plasmid encoding EphA2 or FKBP using Lipofectamine 2000 (Invitrogen) according to the manufacturer’s protocol.

### Cellular treatments with EphrinA1 & western blots

For EphA2 and PKA activity studies, A375 cells were treated with 10 nM EA1-Fc for 10 minutes or 50 nM EA1-Fc for 5 minutes in HEK293T cells transiently transfected with EphA2-GFP. A431 cells were treated with 10 nM EA1-Fc for 1 hour. EA1-Fc was solubilized in DMEM. Control samples received no ligand and were subject to DMEM alone for the treatment time listed above. Cholesterol-modulated samples received ligand stimulation after cholesterol levels were altered. Post-treatment cells were washed twice with warm PBS^++^ (11.9 mM sodium phosphate, 137 mM NaCl, 2.7 mM KCl, 1 mM MgCl2, 0.1 mM CaCl_2_) prior to being lysed.

Cells were harvested and resuspended in ice-cold RIPA buffer supplemented with protease and phosphatase inhibitors (Promega and Sigma Aldrich, respectively). Lysates were centrifuged at 16,200 x g for 20 minutes. Protein concentration was measured with the DC assay (Bio-Rad) and samples were diluted in 4X Laemmli buffer supplemented with dithiothreitol (DTT).

For western blotting, samples were boiled for 5 minutes at 95°C and subject to SDS-PAGE on a 10% gel and proteins were transferred to 0.45 μm nitrocellulose membrane and blocked with 5% BSA in TBS () for 1 hour. The following primary antibodies were used for immunoblotting diluted in 5% bovine serum albumin in TBS and incubated overnight at 4°C: rabbit anti-EphA2 (D4A2) XP (CST 6997; 1:1000), rabbit anti-phospho-EphA2 (Ser897) (D9A1) (CST 6347; 1:1000), rabbit anti-phospho-EphA2 (Tyr588) (D7X2L) (CST 12677;1:1000), rabbit anti-PKA C-α (CST 5842; 1:1000), rabbit anti-phospho-PKA C (Thr197) (D45D3) (CST 5661; 1:1000), rabbit anti-alpha-tubulin (CST 2144; 1:5000), mouse anti-beta-actin (CST 3700; 1:5000). Primary antibodies were detected using host-specific secondary antibodies linked to IRDyes (LI-COR). Western blots were imaged for 680 and 800 nm fluorescence using the Odyssey CLx imaging system (LI-COR). Densitometric analysis of results was carried out using ImageStudioLite software.

### Modulation of cholesterol levels

Methyl-β-cyclodextrin (MβCD) (Acros Organics) was used to remove cholesterol from cultured cells. Cells were incubated for 1 hour with 5 mM MβCD dissolved in DMEM supplemented with 25 mM HEPES, pH 7.4. Zaragozic acid (Sigma Aldrich) was used to remove cholesterol from cultured cells by incubating for 24 hours with 10 μM Zaragozic acid dissolved in DMEM. All treatment incubations were carried out at 37°C with 5% CO_2_.

### Modulation of cAMP levels

Cells were treated with Forskolin (Sigma Aldrich) and Isoproterenol (Thermo Fisher Scientific) for 1 hour at 10 μM and 50 nM, respectively, in DMEM. All treatment incubations were carried out at 37°C with 5% CO_2._

### MTS cytotoxicity

Cells were plated in a clear, flat-bottom 96-well plate to 60% confluency and allowed to adhere for 24 hours. The following day, cells were treated with compounds to modulate cholesterol levels as described above. Post-treatment cells were washed twice with warm PBS^++^ and media was replaced with phenol-free DMEM containing 10% FBS and 100 U/mL penicillin-streptomycin and allowed to grow overnight. Afterwards, MTS reagent (Thermo Fisher Scientific) was added and allowed to incubate at 5% CO_2_ and 37°C for 1.5 hours prior to absorbance being read at 490 nm using a Biotek Cytation V microplate reader with Gen5 software.

### Quantification of cholesterol levels

Cholesterol levels were quantified by the Amplex Red cholesterol assay (Invitrogen) in whole cell lysates prepared from syringe lysis in detergent-free lysis buffer (50 mM Tris-HCl, 250 mM sucrose, 250 μM CaCl_2_, pH = 7.4) following the manufacturers protocol. In brief, the samples were diluted in reaction buffer and an equivalent volume of Amplex Red working solution (300 μM Amplex Red, 2 U/mL cholesterol oxidase, 2 U/mL cholesterol esterase and 2 U/mL horseradish peroxidase) was added. Samples were incubated at 37°C for 1 hour and fluorescence was measured using a BioTek Cytation V microplate reader. Sample excitation occurred in the range of 530-560 nm and emission was detected at ∼590 nm. Cholesterol values were calculated using known cholesterol solutions and normalized to protein content as measured by DC assay (Bio-Rad).

### Lipid extraction for LC-HRMS analysis

Lipid extraction was performed following the procedure detailed by Yang and coworkers^74^. Briefly, A375 cells grown to ∼90% confluency in a 6-well plate were pelleted and washed with PBS. Pellets were then resuspended in 500 µL of ice-cold methanol and vortexed for 5 minutes, followed by the addition of 10 µL of undiluted SPLASH^®^ LIPIDOMIX^®^. The cell suspensions were then freeze-thawed 5x by flash-freezing in liquid nitrogen, warmed to 37 °C in a water bath and briefly vortexed. The cell suspensions were then transferred to a 15 mL conical tube and an additional 500 µL of ice-cold methanol was added, followed by 2 mL of chloroform. The cell suspensions were centrifuged at 1,000 x g for 3 min and the supernatants were transferred to fresh 15 mL conical tubes, where 400 µL of 50 mM citric acid was added followed by an additional 800 µL of chloroform. The solution was vortexed for ∼30 s and then centrifuged at 1,000 x g for 10 min to achieve phase separation. The bottom layer was removed and transferred to a 4 mL glass vial and the solvent was removed under a stream of nitrogen The dried residue was then resuspended in 40 µL of 2:1:1 isopropanol/acetonitrile/H_2_O and transferred to an autosampler vial for LC-HRMS analysis.

### Liquid Chromatography High-Resolution Mass Spectrometry (LC-HRMS) Analysis

Separations were achieved using an Agilent 1290 Infinity UPLC equipped with G4220A binary pump and InfinityLab Poroshell 120 Aq-C18 column (3.0 x 150 mm, 2.7 µm) heated to 40 °C. Injection volume was set to 2 µL. The chromatography gradient was programmed as follows, operating at flow rate of 300 µL/min, where solvent A is acetonitrile:water, 60:40,v/v and solvent B is 10 mM isopropanol:acetonitrile, 90:10, v/v. Solvents A and B are modified with 10 mM ammonium formate and 0.1% formic acid.

| Time (min) | % A | % B |
| --- | --- | --- |
| 5.00 | 57.00 | 46.00 |
| 5.10 | 50.00 | 50.00 |
| 14.00 | 30.00 | 70.00 |
| 21.00 | 1.00 | 99.00 |
| 24.00 | 1.00 | 99.00 |
| 24.10 | 70.00 | 30.00 |
| 28.00 | 70.00 | 30.00 |

Mass spectrometry was performed with an Agilent 6530 Q-ToF mass spectrometer equipped with a Dual AJS electrospray ionization (ESI) source operating in negative polarity with the following parameters:

Gas temperature: 325°C, drying gas flow rate: 7 L/min, nebulizer pressure: 35 psig, sheath gas temperature: 350°C, sheath gas flow: 11 L/min, sprayer voltage: 3.5 kV, nozzle Voltage: 1.0 kV Spectra were acquired in auto MS/MS mode from 100 m/z to 1500 m/z with a scan rate of 500 ms/spectrum in MS^1^ and 125 ms/spectrum in MS^2^ for a total cycle time of 1.225 seconds. Isolation width for precursors was set to narrow (∼1.3 m/z) with a maximum number of precursors per cycle set to 5. Mass correction was performed using purine (112.9855 m/z) and hexakis (1H, 1H, 3H-tetrafluoropropoxy) phosphazine (966.0073 m/z) as reference ions.

Data was analyzed using MS-DIAL^75^ and individual lipid species were identified by matching experimental spectra to reference spectra in the LipidBlast^76^ database. Lipid identification was set with an MS^1^ tolerance of 5 ppm and an MS^2^ tolerance of 10 ppm. Identified lipids were quantitated given the following formula:

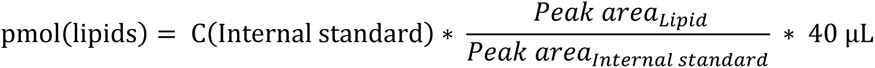

### DIBMALP preparation

C-terminally GFP-tagged proteins (EphA2, FKBP) were expressed in HEK293T cells for 16-18 hours before harvesting the cells. Cell pellets were resuspended in detergent-free lysis buffer (50 mM Tris-HCl, 250 mM sucrose, 250 μM CaCl_2_, pH = 7.4) supplemented with protease and phosphatase inhibitors (Promega and Sigma Aldrich, respectively). Cells were lysed through a series of passages via syringes (25-gauge needle, 20X; vortex 30 seconds; 27-gauge needle, 40X). Supernatants were transferred to fresh Beckman Coulter centrifuge tubes and ultracentrifuged at 100,000xg for 1 hour at 4°C. The pellet was washed with fresh lysis buffer and resuspended via pipetting and 10-20 passes through syringes containing a 25-gauge needle. Resuspended samples were transferred to a fresh Beckman Coulter centrifuge tube and ultracentrifuged at 100,000xg for 1.5 hours at 4°C. The supernatant was removed and resuspended in fresh resuspension buffer (50 mM Tris-HCl, 250 mM NaCl, 90:10 glycerol (v/v), pH = 8.0). Samples were treated with ligand (2uM EA1-Fc Bio-techne; 100nM AP20187 MedChemExpress) for 1 hour on ice prior to being solubilized with 0.15% DIBMA or 1mM DDM overnight at 4°C while shaking. The following day, samples were ultracentrifuged at 100,000xg for 1 hour. Supernatants (DIBMALPs) were collected, and protein concentration was calculated *via* GFP fluorescence and compared to known concentrations of purified GFP (Thermo Fisher Scientific) in a black opaque, flat-bottom 96-well plate using a Biotek Cytation V microplate reader.

### Single-molecule TIRF

DIBMA or DDM solubilized C-terminally GFP-tagged proteins (EphA2, FKBP) were immobilized in a microfluidic chamber prepared on quartz slides coated with a mPEG-silane and 8% biotin PEG-silane mixture (Laysan Bio Inc.) containing two pre-drilled holes. The chamber was prepared by adhering a coverslip to the PEGylated quartz slide with double-sided tape using vacuum grease (Dow Corning) to seal the chamber ends. To immobilize protein samples, 0.02 mg/mL NeutrAvidin protein (Thermo Fisher Scientific) was first incubated in the chamber for 10 minutes followed by the addition of a biotinylated EphA2 or GFP antibody for 20 minutes (Cell Signaling and Rockland Immunochemical Inc., respectively). The microfluidic chamber was rinsed two times with T50 (10 nM Tris-HCl, 50 nM NaCl, pH = 8.0) between each of the above additions. Samples ranging from 100 pM to 5 nM were then added to the chamber and incubated for 30 minutes. After sample incubation, the chamber was rinsed three times with T50 to remove non-specific interactions and a protocatechuic acid/recombinant protocatechuate-3,4-dioxygenase (Sigma Aldrich and Oriental Yeast Co., respectfully) (PCA/rPCD) oxygen scavenging system in Trolox was introduced into the chamber prior to imaging. A customized prism-based TIRF system (as described in prior publications^27,77^) set up with an inverted IX73 Olympus microscope and a customized stage (TIRF Labs Inc.) was used to collect single molecule photobleaching events. Samples were illuminated with a 465nm cable laser (TIRF Labs Inc.). Emission wavelengths were filtered through a custom filter cube (Chroma Technology Corp.) and collected on an EMCCD camera (Andor Technology) using a 100 ms integration time. As described previously (Shushu et al., 2023 and Stefanski et al., 2021), a custom IDL (Harris Geospatial Solutions Inc.) script from the laboratory of Dr. Taekjip Ha (https://github.com/Ha-SingleMoleculeLab/Raw-Data-Analysis) was used to record and extract single molecule trace files. The single molecule traces were analyzed in the Anaconda Navigator Spyder Software with a custom Python code to determine the number of photobleaching steps for each molecule. To account for a 70% maturation efficiency of GFP a custom MATLAB application (see below) was utilized to convert photobleaching steps into percent oligomerization.

### Calculation to convert GFP photobleaching steps into oligomeric distribution (% *n* – mer)

Photobleaching analysis of each intensity trace yields the number of mature GFP only^78,79^. To estimate the total number of GFP, that includes mature and immature ones, we developed a Bayesian method^80–83^. Our method considers stochasticity in the maturation of individual GFP and allows for the isolation of artifacts caused by an efficiency that is less than 100% and biased trace selection caused by traces with only one immature GFP. Specifically, our method models the measured number of mature GFP *w*_n_ based on the total number of GFP *s*_n_ like this

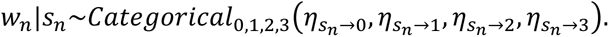

Here, *η*_s→w_ is the probability of having in total *s* = 1,2,3 GFP in a trace selected for analysis with only *w* = 0,1,2,3 measured mature ones. Given that traces with 0 mature GFP are not detected, and so traces with no steps are systematically missed from our data, our probabilities are given by

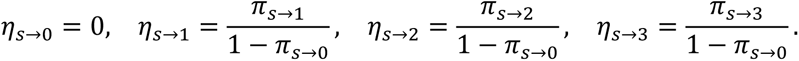

Here, *π*_s→w_ is the probability of having only *w* = 0,1,2,3 mature GFP with *s* = 1,2,3 total ones, irrespective of selecting this trace for analysis or not. In turn, assuming that the maturation of each GFP is independent of the other GFP contributing to the same fluorescence trace, our probabilities are given by

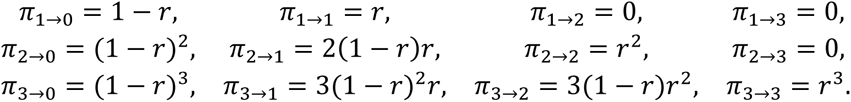

Here, *r* is the maturation efficiency of an individual GFP. Because *s*_n_ is unknown, our method places a Bayesian prior on it of the form

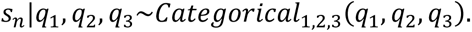

Here, *q*_1_, *q*_2_, *q*_3_ are the probabilities of traces with 1,2,3 GFP, respectively, which is the same as the unknown distribution we seek to estimate. To allow estimation of *q*_1_, *q*_2_, *q*_3_, we apply a non-informative prior of the form

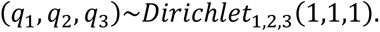

Overall, our model assumes that we measure *N* traces that are indexed with *n* = 1, …, *N*. Using the counts resulting from photobleaching analysis and the model described above, we characterize the posterior probability distribution *p*(*q*_1_, *q*_2_, *q*_3_|*w*_1:N_) via Markov chain Monte Carlo sampling^80,83,84^ and summarize the results by the posterior means and standard deviations of *q*_1_, *q*_2_, *q*_3_.

### Transmission electron microscopy

DIBMALP samples, as described above, were negative-stained for observation by transmission electron microscopy (TEM). For preparation, 5-6 nm thick carbon-coated 200-square mesh copper grids (Electron Microscopy Sciences) were exposed to the sample for 1 minute followed by a 10 second rinse in ddH_2_O. The filter paper was used to adsorb excess liquid from the grids between each step. The grid was then stained with 1% uranyl acetate dissolved in methanol for 1-minute, the excess stain was removed by filter paper, and allowed to air dry overnight. A JEOL JEM 1400-Flash TEM (JEOL USA Inc.) was used to image the prepared grids held in a Fischione 2400 Dual Axis Tomography Holder with an acceleration voltage of 120kV. Images were collected with a Gatan OneView CMOS sensor camera. DIBMALP diameters were measured in ImageJ software.

### Quantification of cAMP levels

HEK293T cells were plated in a clear, flat-bottom 96-well plate to 60% confluency and transduced with the cADDis cAMP sensor according to the manufacturers protocol (Montana Molecular). After transduction, cells were grown overnight at 37°C with 5% CO_2_. The following day, cells were treated with MβCD as described above and control samples (no treatment) received an equivalent volume of DMEM for the duration of MβCD treatment. After treatment, cells were washed 2X with warm PBS^++^ and fluorescence intensity was measured in the same solution using a Biotek Cytation V microplate reader. Samples were excited at 558 nm and emission was detected at 603 nm. Changes in cAMP levels were inferred based on the changes in fluorescence intensity with respect to the control condition.

### Immunostaining and imaging

A375 cells were plated at 80% confluency on a #1.5 glass coverslip, allowed to adhere for 24 hours and then starved overnight. As described above, control samples were treated with fresh DMEM (no treatment) or with/without MβCD and EphrinA1-Fc. Cells were washed with PBS^++^, fixed at 37°C for 15 minutes in 4% paraformaldehyde, and permeabilized for 10 minutes at room temperature with 1% Triton X-100. Cells were blocked with 3% BSA and primary rabbit anti-EphA2 (D4A2) XP (CST 6997; 1:100) antibody was incubated overnight in 1% BSA at 4°C. The following day, cells were washed with PBS^++^ and secondary anti-rabbit conjugated to Alexa-fluor 488 (1:1000) (Thermo Fisher Scientific) was incubated for 1 hour at room temperature. Cells were stained for DAPI (1 μg/mL) (Thermo Fisher Scientific) for 5 minutes at room temperature and mounted to a microscope slide using Diamond Anti-fade mounting media overnight. After curing, cells were imaged using a VT-Hawk 2D array scanning confocal microscope (Visitech intl.) mounted to an IX-83 inverted optical microscope (Olympus). Images were collected using a 60X objective lens (Nikon) in MetaMorph (Molecular Devices).

### Statistical analysis

All statistical comparisons were made using GraphPad software (version 6 for Windows, GraphPad Software, La Jolla California, USA, www.graphpad.com). When only two means were compared, Student’s t-tests were used. When more than two means were compared, one-way analysis of variance (one-way ANOVA) was conducted. If the analysis of variance revealed significant group differences, a Mann Whitney U test was carried out to elucidate the pattern of group differences. When more than two means were compared and two independent variables were present, a two-way analysis of variance (two-way ANOVA) was conducted. If the analysis of variance revealed significant group differences, a Tukey multiple comparison test was used to elucidate the pattern of group differences.

### GPMV preparation and imaging

HeLa cells, transiently transfected with the EphA2-GFP plasmid were grown to 80% confluency in 6-well plates and treated with MβCD as described above. Following treatment, cells were washed 2X with GPMV buffer (10 mM HEPES, 150 mM NaCl, 2 mM CaCl_2_, pH = 7.4) and L_d_ phase marker, DiIC12(3) (Thermo Fisher Scientific), was added to a final concentration of 5 μg/mL in GPMV buffer and incubated for 15 minutes at room temperature, devoid of light. Cells were then washed 2X with GPMV buffer prior to adding freshly-prepared active GPMV buffer containing 25 mM paraformaldehyde and 2 mM DTT. Cells were incubated in active GPMV buffer for 1.5 hours at 37°C. After incubation, GPMVs detached from the cells were gently transferred to a fresh microcentrifuge tube, allowed to settle on ice for 10-45 minutes, and collected by removing ∼20% of the total volume present in the bottom of the tube. 100 μL of the sample was added to a well of a CELLview microscope slide (Greiner bio-one) and allowed to settle for 1.5-2 hours at room temperature devoid of light. Imaging of GPMVs was done using an inverted Zeiss LSM 900/Airyscan laser scanning confocal microscope (ZEISS) with a 63X oil immersion objective. Airyscan images were taken using 1.3X magnification. Images were processed and quantified using ImageJ software.

### Quantification of EphA2-GFP density and probability of co-capture in DIBMALPs

Giant plasma membrane vesicles were generated from HEK293T cells, transiently transfected with EphA2-GFP as described above. Imaging of GPMVs was done using an inverted Zeiss LSM 900 confocal microscope (ZEISS) with a 63X oil immersion objective. Confocal images were taken with the pinhole set to 30 μm. Images across conditions were acquired using the same laser power and camera gain settings. Image processing and quantification were done using ImageJ software. Using the circle function on ImageJ, the raw intensity of EphA2-GFP within each vesicle was recorded. Additionally, the line function on ImageJ was used to obtain the diameter of each GPMV. The surface area of each GPMV was calculated using the diameter and pinhole size and assigned a GFP intensity value. The fluorescence of the GPMV samples were compared against the fluorescence intensity of purified GFP at a known concentration (100 nM) and buffer blanks were subtracted. This concentration was converted to estimate the number of EphA2-GFP molecules per micron on the GPMV surface. Using the median density of EphA2-GFP molecules per square micron (across all replicates) and multiplying this by a range of DIBMALP sizes provided the probability of randomly co-capturing EphA2-GFP monomers in a single DIBMALP.

### Modeling of the TM-JM peptides

The NMR structure of EphA2 TM dimer (PDB ID: 2K9Y)^48^ was obtained from www.rcsb.org. For modeling of the TM-JM peptide [E^530^GSGNLAV<u>IGGVAVGVVLLLVLAGVGFF</u>IHRRRKNQRAR^568^], we extracted the TM region and part of the N- and C-terminal residues of EphA2 (E^530^–K^563^) from the NMR structure and then the remaining C-terminal residues from N^564^–R^568^ were modeled as an extended conformation of amino acids (ϕ, ψ= ±120°) in PyMOL (The PyMOL Molecular Graphics System, Version 2.5. Schrödinger, LLC).

### Set up for Coarse-Grain (CG) molecular dynamics simulation

To analyze the dimerization of TM-JM peptides, the monomers were positioned 50 Å apart from each other. Subsequently, the atomistic (AT) models of TM-JM peptides were transformed into a coarse-grained (CG) representation using the martinize2.py workflow module from the MARTINI 3 force field,^85^ considering the secondary structure assignments from DSSP.^86^ We employed an elastic network to enhance the stability of the helical secondary structure in the TM monomers. We used default values of the force constant of 500 kJ/mol/nm^2^ with the lower and upper elastic bond cut-off to 0.5 and 0.9 nm respectively. CG simulations were performed using GROMACS version 2016.5.64.^87^ Next, the peptides were introduced, positioned perpendicular to the membrane. We constructed two distinct systems, differentiating solely based on lipid composition: the first system comprised POPC (55%), Cholesterol (40%), and PIP2 (5%), while the second had a similar setup but without Cholesterol, consisting of POPC (95%) and PIP2 (5%). We utilized the insane.py^88^ script to establish the lipid bilayer. For system 1, this typically included 175 POPC, 127 CHOL, 15 PIP2 lipids, and 4470 CG water molecules. For system 2, the setup consisted of 303 POPC, 15 PIP2, and 4213 CG water molecules. The systems were encased in a cubic box measuring 100 × 100 × 100 Å^3^. The pH of the systems was considered neutral. All the simulations were run in the presence of regular MARTINI water and were neutralized to 0.15M NaCl. The systems were equilibrated for 500 ps. The long-range electrostatic interactions were used with a reaction type field having a cutoff value of 11 Å. We used potential-shift-verlet for the Lennard-Jones interactions with a value of 11Å for the cutoff scheme and the V-rescale thermostat with a reference temperature of 320 K in combination with a Berendsen barostat with a coupling constant of 1.0 ps, compressibility of 3 × 10^-4^ bar^-1^, and a reference pressure of 1 bar was used. The integration time step was 20 fs. All the simulations were run in quadruplicate for 4 µs each.

### Simulation Data Analysis

Trajectory analysis was conducted using the integrated modules within GROMACS. Contact maps depicting the TM regions and with the lipids were generated, employing a cutoff of 6 Å for both backbone and side-chain atoms. Subsequently, the data were visualized and plotted using GraphPad Prism (version 6 for Windows, GraphPad Software, La Jolla California, USA, www.graphpad.com).

### SDS-PAGE

Lipid/peptide films were prepared as above with the TMJM63 peptide at a lipid:peptide ratio of 300:1. Films were resuspended in 19.3mM HEPES, 1mM EGTA and shaken at room temperature for 3 hours to allow disulfide bond formation. After MLV formation, SDS buffer was added to a final concentration of 150mM along with sample buffer with/without DTT. Samples were boiled for 5 minutes at 95°C, ran on a 16% tricine gel and stained using a Pierce Silver Stain Kit (Thermo Fisher Scientific). Densitometry was performed using ImageStudioLite software.

## Supporting information

Supplementary figures 1-15, table 1, references

## Acknowledgements

This work was supported by NIH grants R35GM140846 (F.N.B) and R35GM142946 (R.L.). We are grateful to Amit Joshi for the use of his Zeiss confocal microscope.

