## Supplementary figures 1-15, table 1, references for "Cholesterol inhibits assembly and activation of the EphA2 receptor"

**Supporting Information for**
### Cholesterol inhibits oncogenic EphA2 activation

Ryan J Schuck, Alyssa Ward, Amita R Sahoo, Jennifer A Rybak, Robert J Pyron, Thomas N Trybala, Timothy B Simmons, Joshua A Baccile, Ioannis Sgouralis, Matthias Buck, Rajan Lamichhane, and Francisco N Barrera

**This PDF file includes:**

Figures S1 to S15
Tables S1
SI References

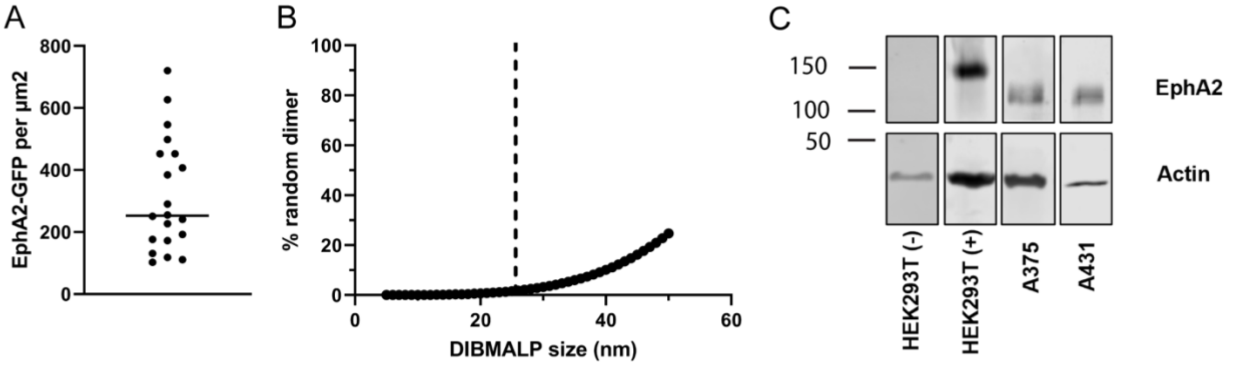


**Supplementary Figure 1: EphA2-GFP membrane density and probability of random co-capture in DIBMALPs.** (A) Quantification of EphA2-GFP membrane density. Since data is not normally distributed, we report the median density = 253 EphA2 molecules per $\mu m^2$. (B) Probability of two EphA2-GFP molecules randomly captured in a DIBMALP as a function of DIBMALP size at 253 EphA2-GFP per $\mu m^2$. Dashed line represents average DIBMALP size (25 nm). (C) HEK293T cells do not express significant levels of endogenous EphA2. Representative western blot using anti-EphA2 antibody for non-transfected (-), and EphA2-GFP transfected (+) HEK293T cells. Endogenous EphA2 is observed in A375 and A431 cells. EphA2-GFP = 152 kDa, and EphA2 = 125 kDa.

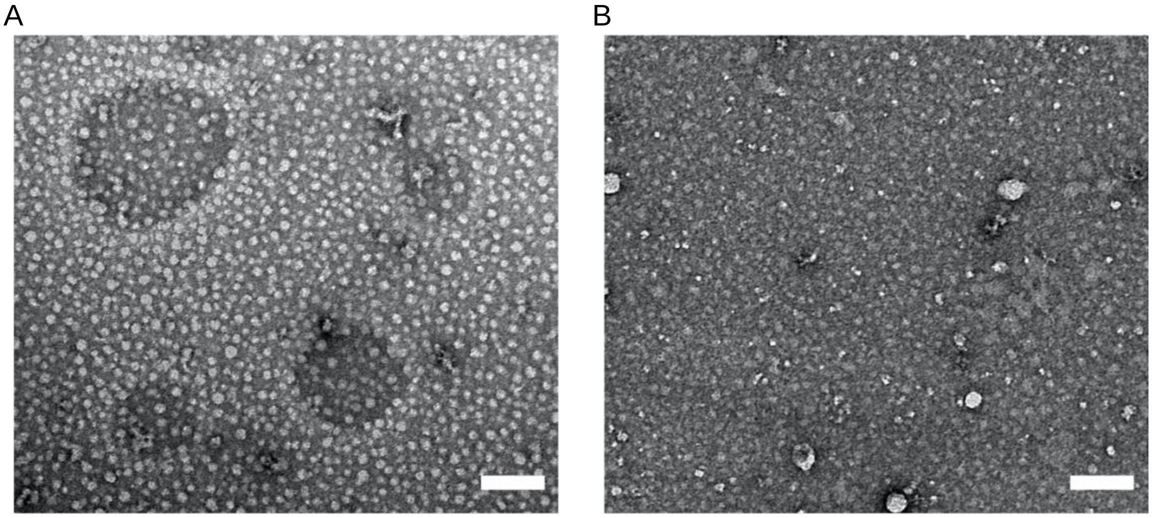


**Supplementary Figure 2: TEM images of DIBMALPs and size distributions.** (A) Representative TEM image of DIBMALPs formed under control conditions. (B) Representative TEM image of DIBMALPs formed following MβCD treatment.  Scale bar is equal to 100 nm.

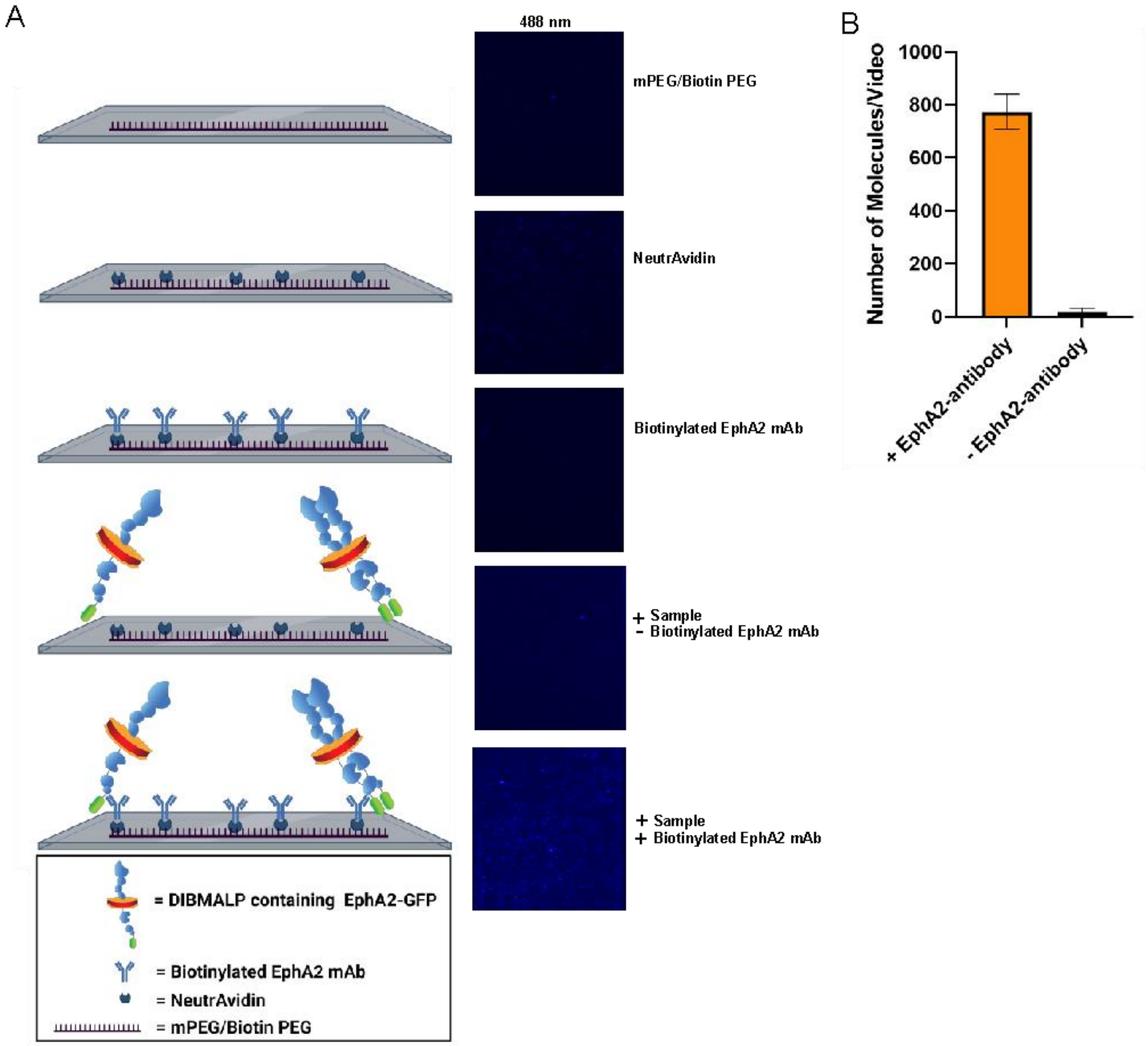


**Supplementary Figure 3: Immobilization of DIBMALPs containing EphA2-GFP on microscope slides.** (A) Steps of slide coating and sample isolation are shown in the left column, with representative TIRF images under 488 nm excitation shown in the right column. (B) Number of molecules in the 488 nm channel per video in the presence and absence of EphA2 antibody. Mean ± S.D. for 3-5 videos.

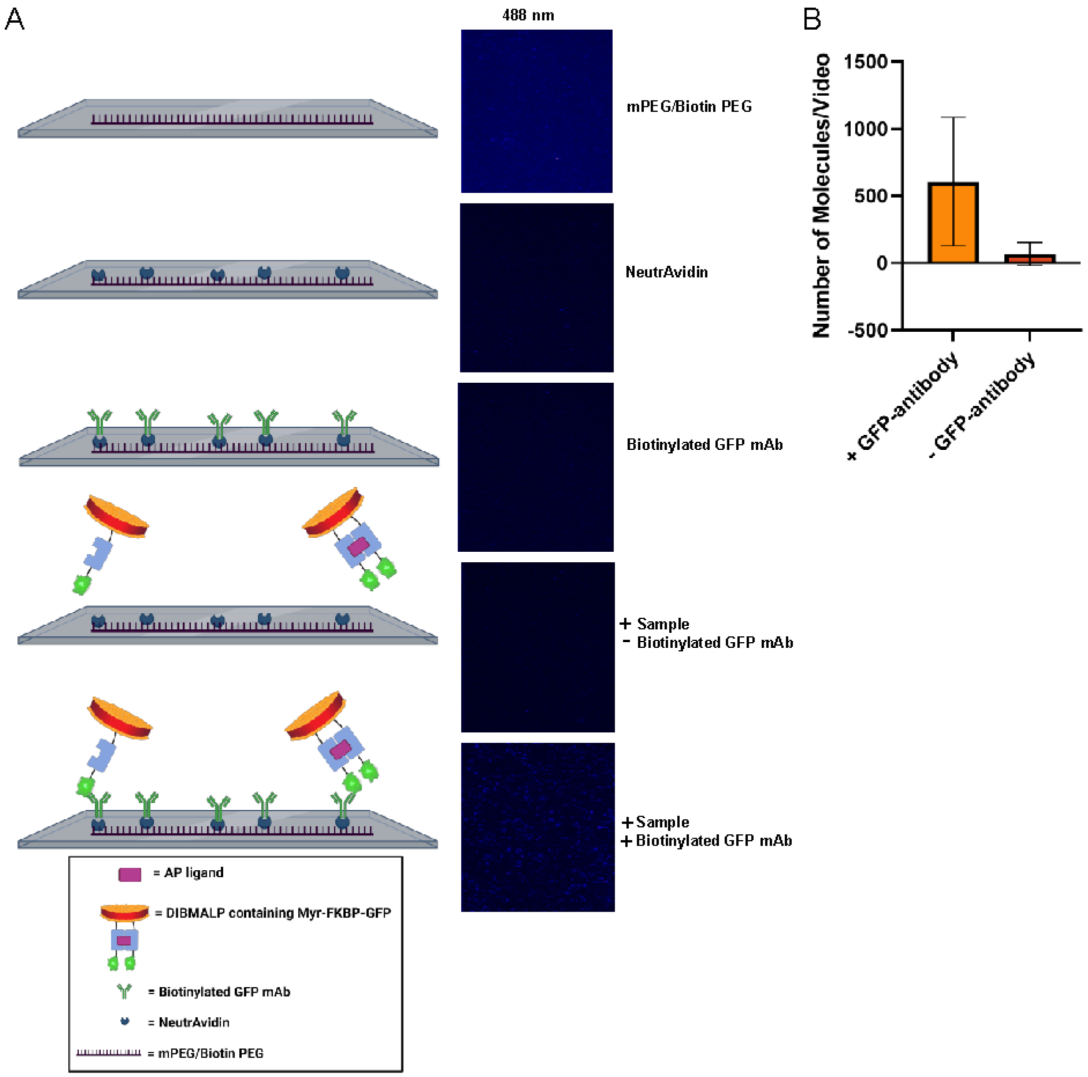


**Supplementary Figure 4: Immobilization of DIBMALPs containing FKBP-GFP on microscope slides.** (A) Steps of slide coating and sample isolation are shown in the left column, with representative TIRF images under 488 nm excitation are shown in the right column. (B) Number of molecules in the 488 nm channel per video in the presence and absence of GFP antibody. Mean ± S.D. for 3-5 videos.

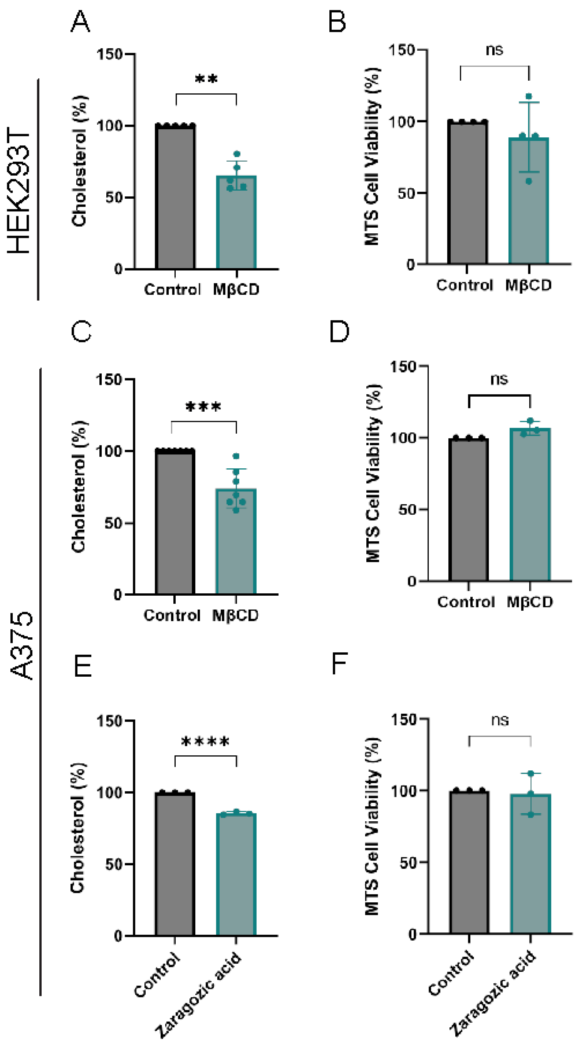


**Supplementary Figure 5: MβCD treatment reduces cholesterol content without impacting cell viability**. (A) Cholesterol quantification and (B) cell viability in HEK293T cells following treatment with MβCD. (C) Cholesterol quantification and (D) cell viability in A375 cells following treatment with MβCD. (E) Cholesterol quantification and (F) cell viability in A375 cells following treatment with zaragozic acid. Data are normalized to control conditions and represented as mean ±S.D., *p*-values from unpaired t test; **, $p < 0.01$; ***, $p < 0.001$; and ****, $p < 0.0001$.

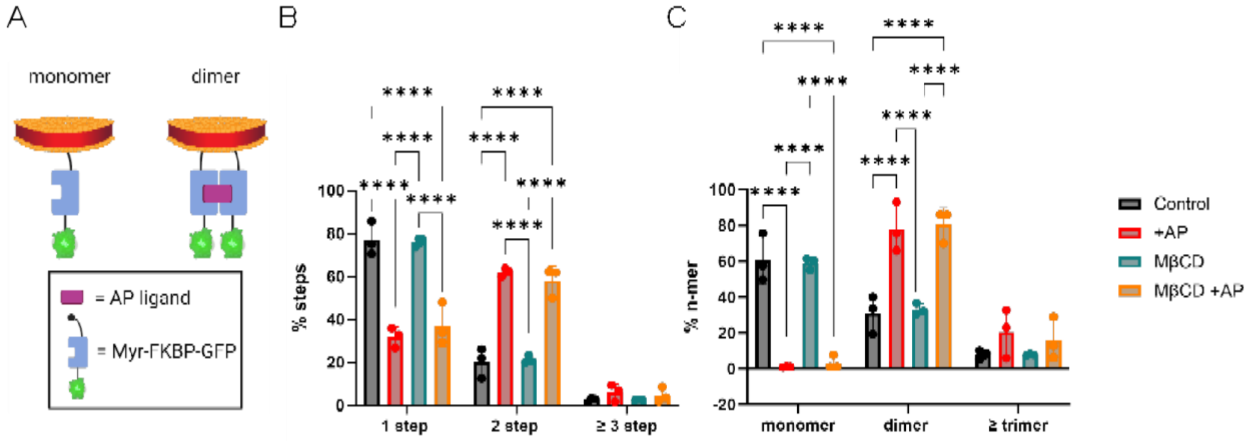


**Supplementary Figure 6: Cholesterol reduction does not impact FKBP dimerization.** (A) Schematic representation of DIBMALPs containing Myr-FKBP-GFP (monomer, left). FKBP-GFP dimerization is induced using the AP ligand (dimer, right). (B) Experimental step distribution of FKBP-GFP photobleaching in control conditions (black), in the presence of AP (red), after MβCD treatment (teal) or with both treatments (orange). (C) Oligomeric distribution of experimental FKBP-GFP photobleaching data obtained from panel B after GFP maturation correction. *p*-values are from two-way ANOVA followed by Tukey multiple comparison test. ****, $p < 0.0001$.

A

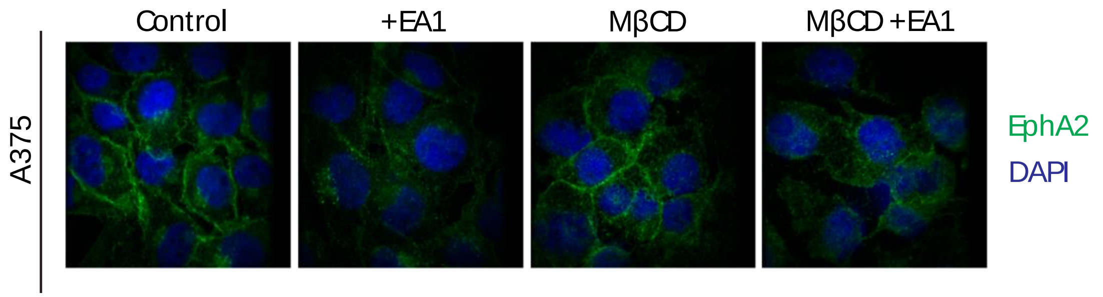


**Supplementary Figure 7: Cellular distribution of EphA2.** (A) Immunofluorescence analysis of EphA2 in the presence and absence of EA1. Images were collected in control conditions or following MβCD treatment. Experiments were performed in A375 cells expressing endogenous EphA2 (green). Nuclei were stained with DAPI (blue).

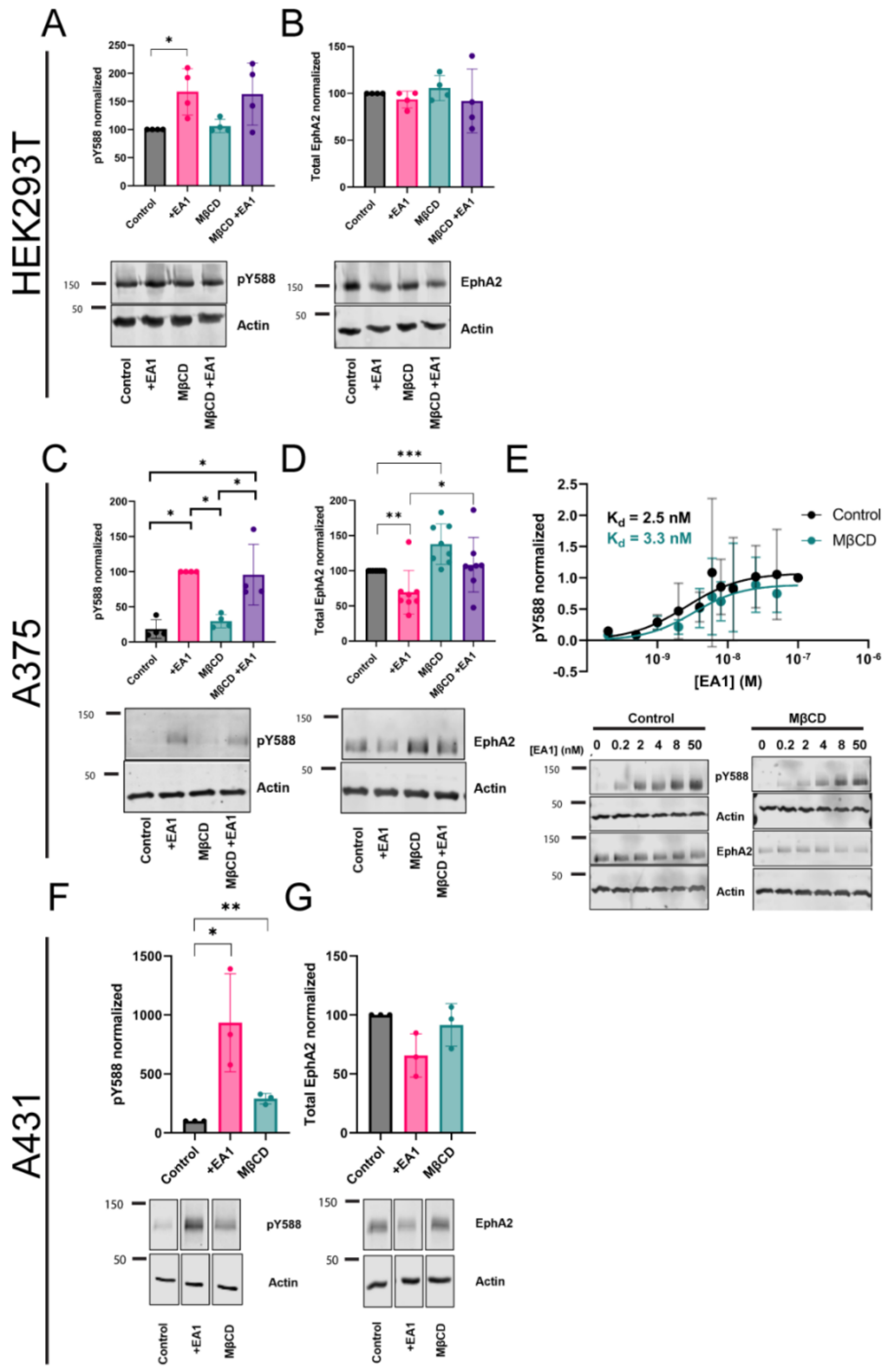


**Supplementary Figure 8: Effect of Chol extraction in pTyr and total EphA2 levels.** Data are shown for HEK293T (A-B), A375 (C-E) and A431 (F-G) cells. We show quantification (mean ±S.D.) of pY588 (A, C and F) and total EphA2 levels normalized with to actin (B, D and G). Panel E shows dose response curves of EphA2 pY588 following treatment with varying concentrations of EA1. Data were fit to the Hill equation to determine the EA1 dissociation constant (control $K_d = 2.5 \pm 1.5$ nM; MβCD $K_d = 3.3 \pm 1.3$ nM). Quantitative comparisons between treatments were normalized to control conditions. *p*-values are from one-way ANOVA followed by Mann-Whitney *U* test or *t* test. *, $p < 0.05$; **, $p < 0.01$;***, $p < 0.005$.

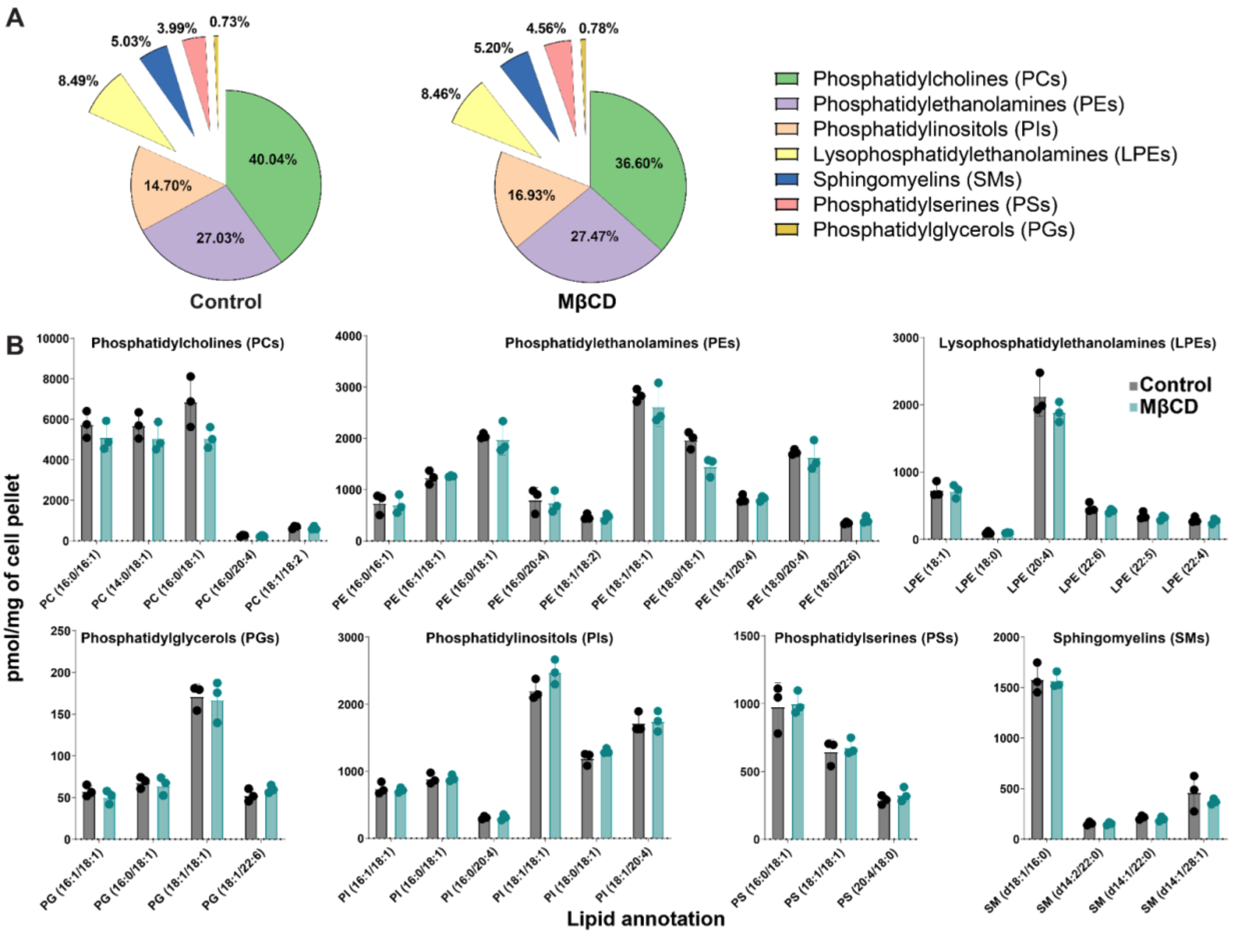


**Supplementary Figure 9: Methyl-β-cyclodextrin does not significantly alter the phospholipid composition of A375 cells.** (A) Relative abundances of identified lipids expressed as mol percentages normalized to 1 mg of A375 cell pellet. (B) Normalized amounts (n=3) of identified lipids sorted by individual class. Statistical significance between control and MβCD treated samples was tested with Welch's t-test and no statistical significance was observed (α = 0.05) for any individual lipid. Identified lipids are annotated following the abbreviation conventions used by LIPIDMAPS[1].

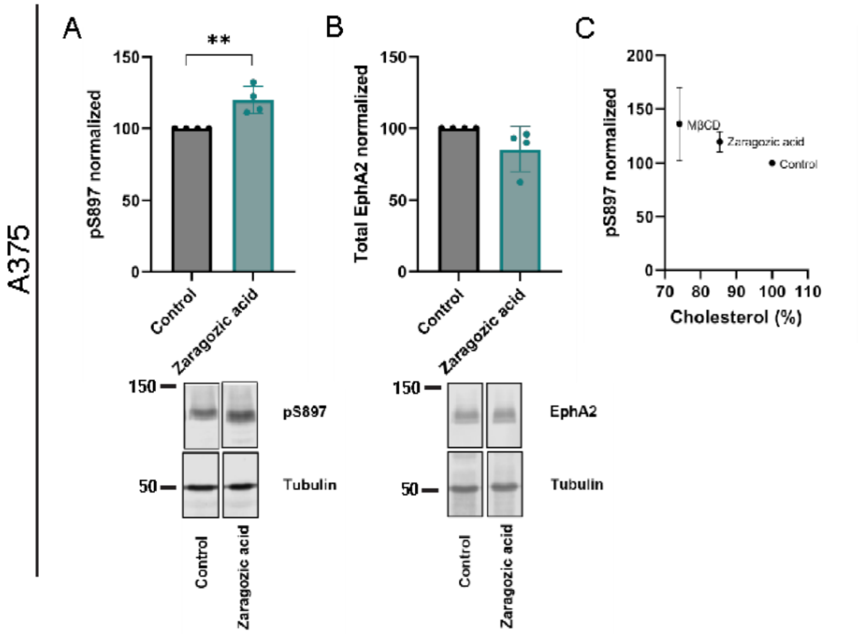


**Supplementary Figure 10: Zaragozic acid treatment causes a moderate Chol reduction that increases pS897 EphA2 in A375 cells**. Effect of zaragozic acid treatment on pS897 (A) and total EphA2 (B). Data shown in panel A is normalized to respective total EphA2. Total EphA2 signal in panel B is normalized with respect to tubulin. (C) Comparison of the effect of MβCD and zaragozic acid. Data shown panels A-C are represented as mean ± S.D., *p*-value in panel A is from unpaired t-test; ** $p \leq 0.01$. Statistics in panel B not significant.

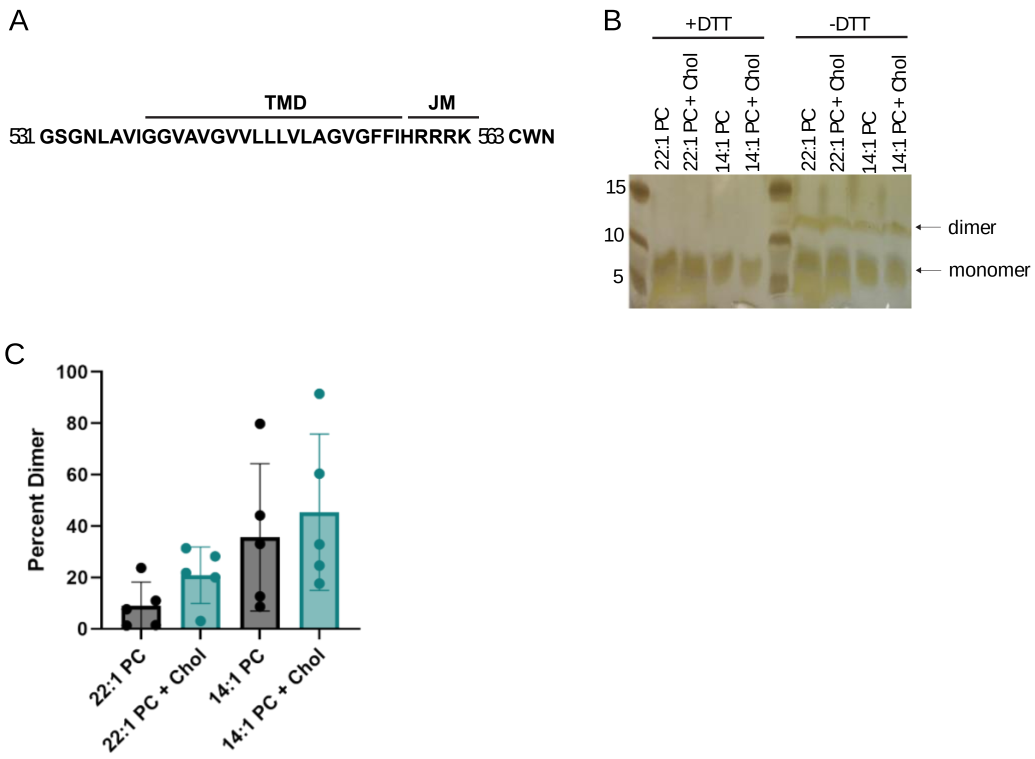


**Supplementary Figure 11: Cholesterol does not influence EphA2-TM dimerization.** (A) TMJM peptide sequence, comprised of EphA2 residues 531-563 with CWN residues added at the C-terminus. (B) Representative SDS-PAGE of TMJM in 14:1 and 22:1 PC vesicles in the presence and absence of 40% cholesterol. Addition of 5 mM DTT eliminates disulfide-mediated dimer bands (left). Monomer and disulfide-mediated dimers can be seen in non-reducing conditions (right). (C) Quantification of bands in panel B. Statistics in panel C show no significance.

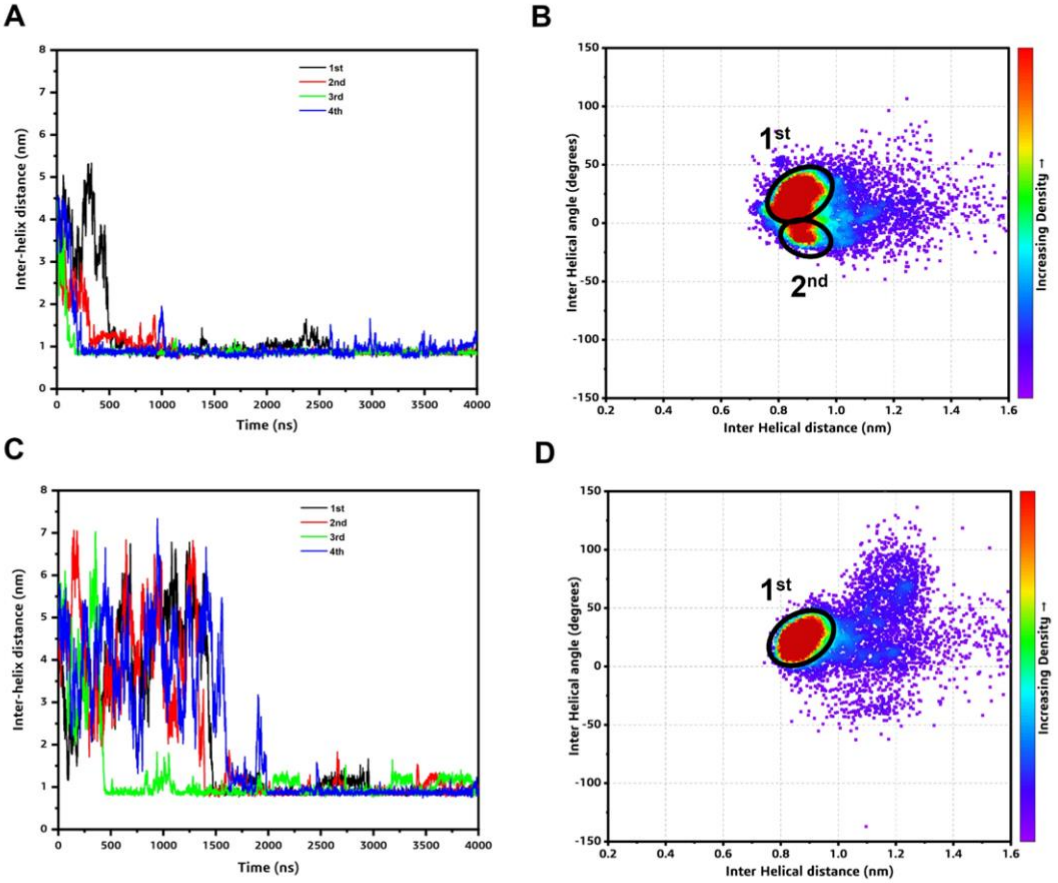


**Supplementary Figure 12: Comparison of the association of the EphA2 TM regions in MD experiments.** Inter-helical distance (COM) plots showing the association between the TM regions of EphA2 in the presence (A) and absence of cholesterol (C) in the membrane. 1st, 2nd, 3rd, and 4th simulation results are shown in black, red, green, and blue lines, respectively. Right panel- 2D distribution plot (interhelix angle vs. distance) between the EphA2 TMs in the presence (B) and absence of cholesterol (D) in the membrane. One population cluster is observed in all cases ($1^{st}$) with a minor shoulder observed in the presence of Chol ($2^{nd}$).

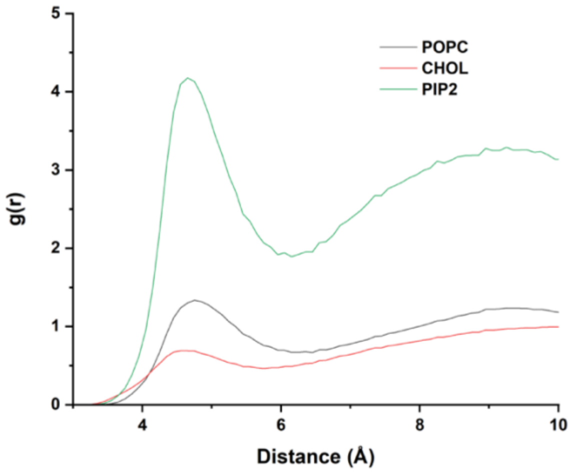


**Supplementary Figure 13: Comparison of the radial distribution functions -g(r)- of lipids around EphA2 TM.**

| | Clusters | X (degrees) | RMSD (Å) |
|---|---|---|---|
| **EphA2 TM (NMR)** | | 15 | |
| **EphA2 TM with Chol** | $1^{st}$ | 25.5 ± 8.5 | 3.8 ± 0.1 |
| | $2^{nd}$ | -10.5 ± 5.0 | 3.8 ± 0.5 |
| **EphA2 TM without Chol** | $1^{st}$ | 26 ± 11 | 5.4 ± 0.2 |

**Supplementary Table 1: Comparison of CG-simulated EphA2 TM dimers of both the systems with the NMR structures.** Five representatives from each population clusters are considered for calculation.

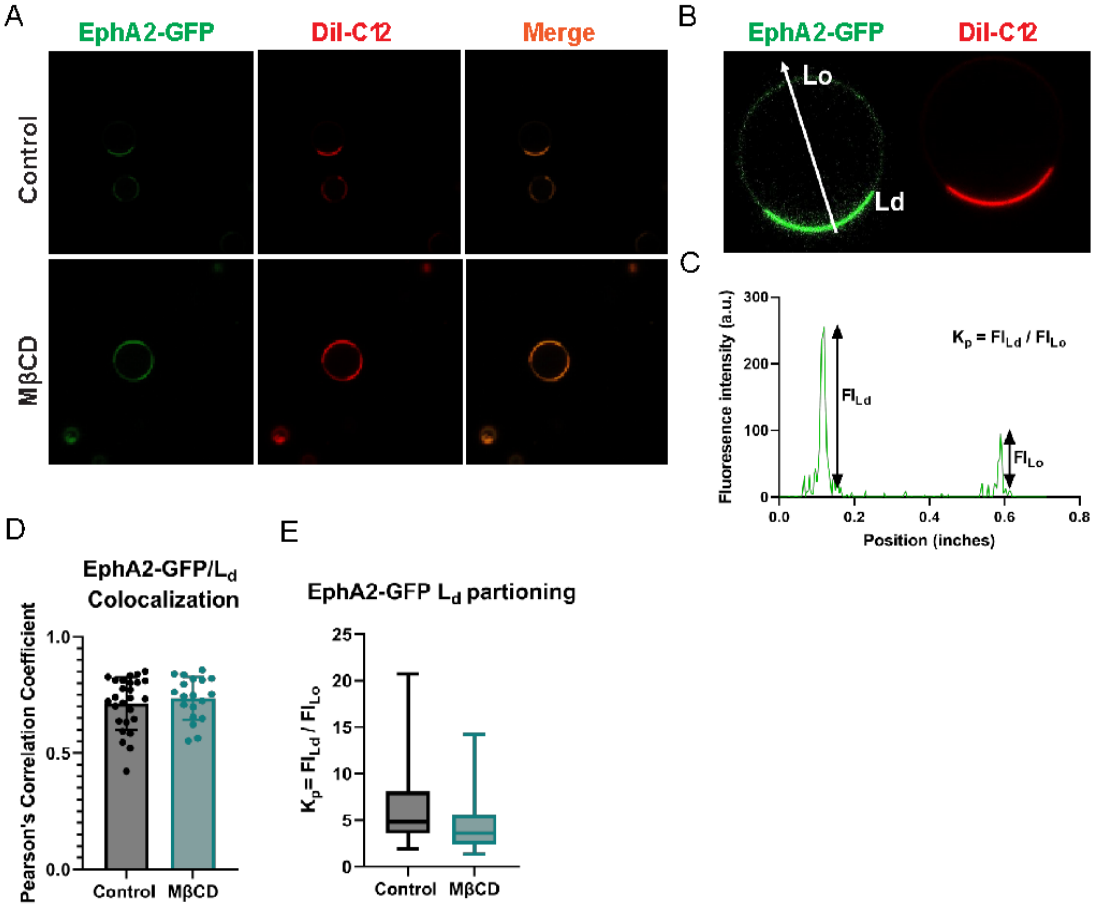


**Supplementary Figure 14: Cholesterol reduction does not affect EphA2 localization to cholesterol rich domains in GPMVs.** (A) Representative confocal images of EphA2 lipid phase partition after MβCD treatment. GPMVs were formed in HeLa cells transiently transfected with EphA2-GFP (green). The dye DiI-$C_{12}$ marking the $L_d$ phase is shown in red. (B) The white arrow represents the line used for the fluorescence intensity scan between colocalized EphA2-GFP/ DiI-$C_{12}$ signal. (C) The ratio of GFP fluorescence in the $L_d$ and $L_o$ phase allows to calculate the partition coefficient ($K_p$) of $L_d$ phase preference of EphA2-GFP. (D) Quantitative colocalization (Pearson's) between EphA2-GFP and DiI-$C_{12}$ signal under control conditions or after MβCD treatment. (E) Partition coefficient of EphA2-GFP. Statistics in panels D & E show no significant changes.

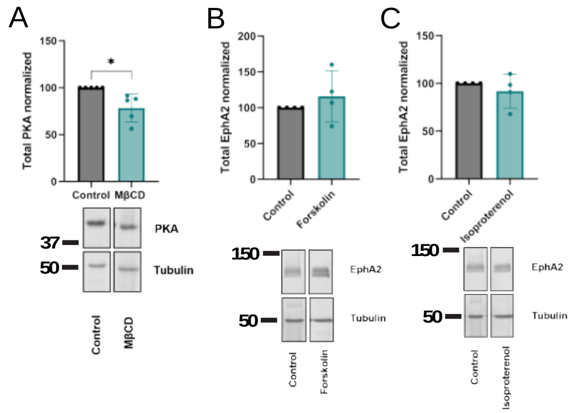


**Supplementary Figure 15: Cholesterol reduction increases cAMP levels and PKA activity.** (A) Quantification of total PKA following treatment with MβCD in A375 cells. (B-C) Quantification of total EphA2 following treatment with forskolin and isoproterenol, respectively. The total PKA signal in panel A is normalized with respect to tubulin. The total EphA2 signals in panel B-C are normalized with respect to tubulin. Quantitative comparisons between treatments were made with respect to normalized control conditions. Bar graphs show mean ± S.D., *p*-values in panels A-C not significant.
